# Dopaminergic Neurons Linking Threat Processing to Cardiac Modulation and Locomotor Responses

**DOI:** 10.64898/2025.12.21.695849

**Authors:** Masato Tsuji, Daisuke Jinkoma, Yuki Uemura, Ayako Ogasawara, Kazuo Emoto

## Abstract

Heartbeat and behavior are tightly coordinated during defensive states, yet the neuronal mechanisms linking threat processing to cardiac modulation—and the potential contribution of cardiac dynamics to behavioral output—remain poorly understood. Here we show in *Drosophila* that mechanical threat triggers locomotion together with cardiac deceleration. We identify two pairs of dopaminergic neurons, termed DA-WED neurons, that mediate this cardiac response: silencing these neurons markedly attenuates threat-induced cardiac deceleration, whereas optogenetic activation induces cardiac deceleration in the absence of threat. Calcium imaging further shows that DA-WED neurons are activated by mechanical threat. Linking cardiac dynamics to behavior, direct optogenetic manipulation of cardiomyocytes that quantitatively reproduces threat-evoked cardiac deceleration is accompanied by increased locomotion. Together, these results identify a dopaminergic descending pathway that links threat processing to cardiac modulation and suggest that cardiac dynamics may contribute to shaping defensive behavior.

## INTRODUCTION

Heartbeat dynamics are tightly coordinated with behavioral state. Defensive states, for example, are accompanied by coordinated modulation of cardiac activity, including transient cardiac deceleration during threat detection and cardiac acceleration during active coping in mammals^1^. Such tight coordination between behavioral state and cardiac dynamics is likely advantageous for survival: cardiac deceleration may minimize the brain-wide representations of the heartbeat^2–5^ to sharpen sensory processing^16^, whereas cardiac acceleration boosts the delivery of metabolic resources to muscles^7^. Although much of the existing work on heart–behavior coupling has focused on mammals, comparable threat-associated cardiac modulation has also been reported in invertebrates. In Drosophila and decapod crustaceans, for instance, sensory stimuli elicit time-locked heart rate changes that parallel defensive behaviors, suggesting that coordinated heart–behavior responses are evolutionarily widespread ^8–10^. Despite this well-documented coupling, the circuit mechanisms linking threat processing in the brain to cardiac modulation—and whether cardiac dynamics contribute to behavioral output—remain incompletely understood.

In mammals, heartbeat is governed by sympathetic and parasympathetic branches of the autonomic nervous system (ANS) that innervate the heart^7,11–18^. Sympathetic output that accelerates the heartbeat arises from brainstem and hypothalamic circuits and projects through spinal intermediates to the heart^7^. On the other hand, the parasympathetic nervous system, which decelerates the heartbeat^7,19^, originates from the dorsal motor nucleus of the vagus nerve and nucleus ambiguus, and projects directly to the heart^19,20^. These autonomic pathways are engaged in threatened animals by the so-called fear circuitry in the brain^21–23^ to shape cardiac responses, typically producing cardiac deceleration followed by acceleration^1,24–29^. Circuit-wise, the periaqueductal gray (PAG) is considered the output hub of the fear circuitry and modulates the heartbeat through the hypothalamus, midbrain, pons, and medulla oblongata^30^. In addition, hypothalamic nuclei such as the dorsomedial hypothalamus (DMH)^31^ and the arcuate nucleus (ARC), as well as the anterior cingulate cortex (ACC)^32^, also modulate the heartbeat upon threats^33–39^.

In parallel, ascending feedback from the heart to the brain has attracted increasing attention. Such interoceptive signaling^40^ is conveyed by sensory neurons innervating the heart and aortic arch^40^, which project to NTS and onward to hypothalamic and cortical regions including paraventricular nucleus of the hypothalamus (PVN) ^41^, ACC, insular, and prefrontal cortex^41^. These pathways have been implicated in learned fear^42^, anxiety-like behavior^43^ or sleep^44^. Together, these findings suggest that bidirectional communication between brain and heart may shape physiological and behavioral responses during defensive states. However, how central circuits orchestrate threat-associated cardiac modulation, and whether cardiac dynamics influence defensive behavior, remain unresolved.

The fruit fly *Drosophila melanogaster* provides a genetically tractable system to dissect brain–body interactions. The *Drosophila* heart, an evolutionarily conserved organ^45–48^, consists of two rows of cardiomyocytes forming a tubular structure that runs through the midline of the abdomen^49,50^. Like mammalian heart, *Drosophila* heart receives rich neuronal projections throughout^51,52^. Recent work has shown that visual threat induces cardiac deceleration in *Drosophila*^10^, suggesting the existence of the neuronal mechanism in the brain that modulates the heart. However, the neuronal substrate of such a mechanism, and the behavioral consequences of cardiac modulation during threat, remain largely unknown.

We have recently reported that a train of air puffs elicit hyperactivity, avoidance of puffs, and aversion from an otherwise neutral visual object, suggesting that air puffs serve as mechanical threat^53^. Building on this work, we established an experimental paradigm to assess how mechanical threat modulates heartbeat and behavior. Using this system, we identify two pairs of dopaminergic neurons in the brain, termed DA-WED neurons, that are responsible for threat-induced cardiac deceleration. We further examine how cardiac deceleration relates to threat-induced locomotion. Together, our findings identify two pairs of dopaminergic neurons that link threat processing in the brain to cardiac modulation and suggest that cardiac dynamics may contribute to shaping defensive behavior.

## RESULTS

### I. Air puffs induce cardiac deceleration and locomotion

We have previously reported that a train of air puffs serve as mechanical threats, eliciting a host of fear-like behaviors such as hyperactivity, avoidance of puffs, and aversion from an otherwise neutral visual object^53^. This motivated us to test how the air puff might influence heartbeat, and its relation to behavior. For this purpose, we developed a paradigm to monitor heartbeat and behavior simultaneously (Fig. 1a). Our paradigm images the heart chamber closest to the thorax, called conical chamber, which is visualized through the dorsal cuticle by cuticle-penetrating infare-red light (Fig. 1a, “CAM1”). Simultaneously, the fly’s behavior is recorded from the side of the fly (Fig. 1a, “CAM2”). We further developed a set of software to automatically detect heartbeat and behavior (Supplementary Movie 1). Our heartbeat detection software takes two consecutive frames and calculates for each pixel the optical flow, which are used to quantify the overall relaxation (diastole) or shrinkage (systole) of the heart (Supplementary Fig. 1a-c). Based on such quantification, our software detects the timings at which the heart was shrunk most (systole peak), which were counted to obtain the heart rate (HR). Our analysis pipeline for behavior detection first tracks the coordinate of body parts (joints of each leg, eye, neck, haltere, rostral tip of the abdomen, all on the right side of the fly), utilizing SLEAP software^54^. The tracked coordinates are then fed into our custom support vector machine-based program to detect locomotion. Our analysis pipeline allowed us to visualize the HR and locomotion over time in a synchronized fashion.

**Fig. 1.**
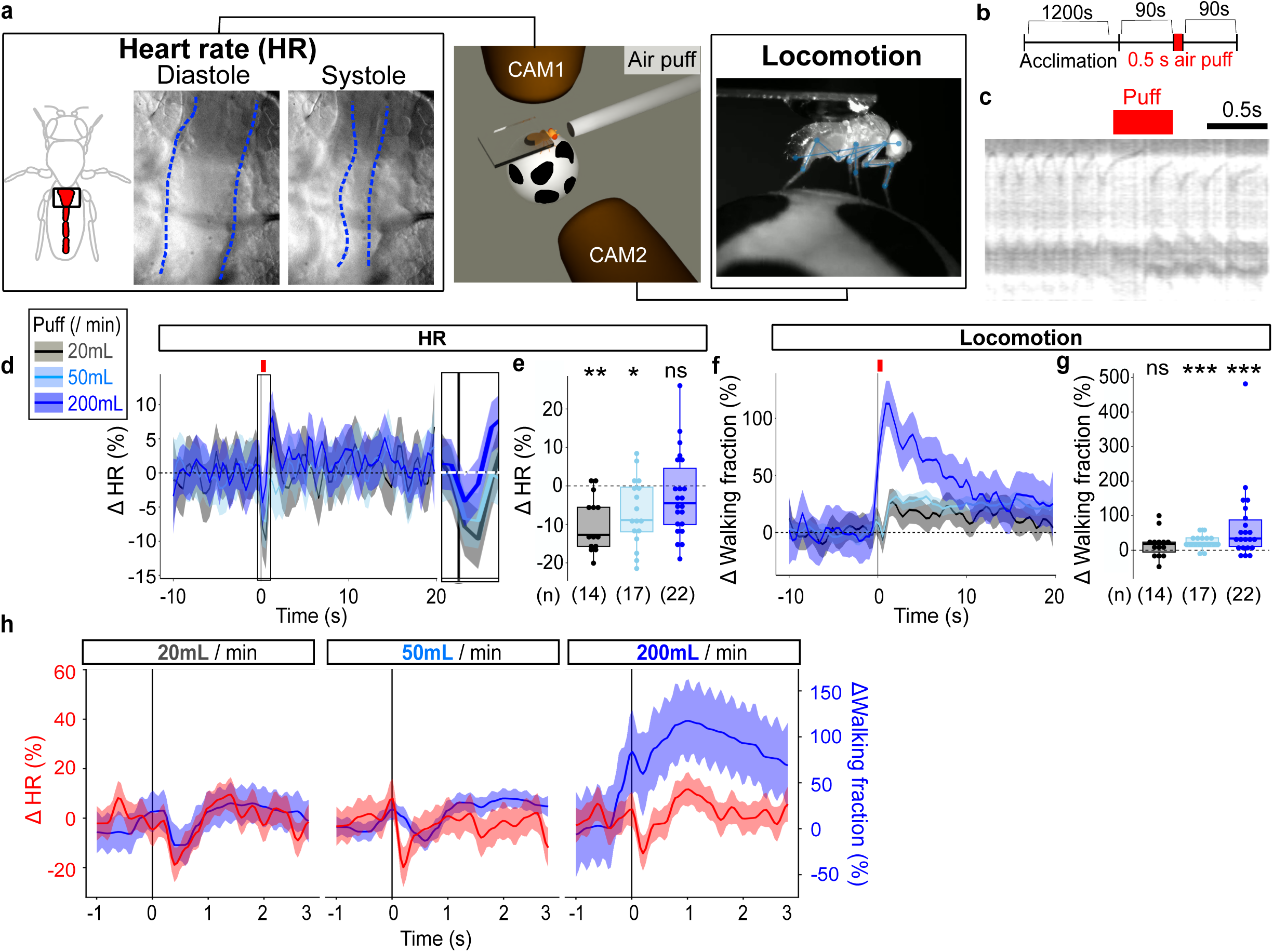
Mechanical threat induces cardiac deceleration and locomotion. **a.** Schematics of the paradigm. The fly is fixed on an air-supported ball. Left: the heart rate (HR) was videotaped by an IR-sensitive camera from the dorsal side through the cuticle (“CAM1”). Of the four cardiac chambers, our system videotaped the chamber closest to the thorax called conical chamber. We could easily distinguish diastole and systole in the captured video (blue dashed lines indicate the heart wall). Right: simultaneously, locomotion was videotaped from the right side of the fly (“CAM2”). Blue dots and lines indicate automatically tracked body parts. **b.** The time course of the experiment. An air puffs of 0.5 s was applied once. The trial was repeated 10 times for each fly, the results of which were averaged. **c.** An example M-mode of the heartbeat. An air puff was applied during the red-shaded time window. **d.** Time course of the HR (% change from baseline defined as the average during the 10 s preceding puff onset). N = 14, 17, 22 for 20mL/min (gray), 50mL/min (sky blue), and 200mL/min (dark blue), respectively. The inlet shows the magnified view of the cardiac deceleration indicated by the box. **e.** Change in HR, calculated for each fly as the % difference between the mean fraction during the 1 s following stimulation onset and the mean fraction during the 10 s baseline period preceding stimulation. N = 14, 17, 22 for 20mL/min (gray), 50mL/min (sky blue), and 200mL/min (dark blue), respectively. **p < 0.01, *p < 0.05, n.s.: p > 0.05, Wilcoxon’s signed rank test followed by Bonfferoni correction. **f.** Time course of the walking fraction (% change from baseline defined as the mean fraction during the 10 s preceding puff onset). N = 14, 17, 22 for 20mL/min (gray), 50mL/min (sky blue), and 200mL/min (dark blue), respectively. ***p < 0.001, n.s.: p > 0.05, Wilcoxon’s signed rank test followed by Bonfferoni correction. **g.** Change in walking fraction, calculated for each fly as the % difference between the mean fraction during the 10 s following stimulation onset and the mean fraction during the 10 s baseline period preceding stimulation. N = 14, 17, 22 for 20mL/min (gray), 50mL/min (sky blue), and 200mL/ min (dark blue), respectively. ***p < 0.001, n.s.: p > 0.05, Wilcoxon’s signed rank test followed by Bonfferoni correction. **h.** Changes in HR and walking fraction jointly plotted for -1 to 3s around the stimulus onset, for each puff strength. N = 14, 17, 22 for 20, 50, and 200mL/min, respectively.

Using our paradigm, we tested how the air puff might influence the HR and locomotion. Specifically, we applied for each fly a 0.5 s air puff (either 20 mL/min, 50 mL/min, 200 mL/min). We repeated 10 trials for each fly with an inter-puff interval of 90s (Fig. 1b). The results of 10 trials were averaged, as we did not observe noticeable trial effects (Supplementary Fig. 2a, c). We found that the air puff transiently decreased the HR (Fig. 1c-e; Supplementary Fig. 2a, b; Supplementary Movie 1). A similar tendency was observed but did not reach statistical significance for the strongest puff (200 mL/min), which is likely due to the following cardiac acceleration (Fig. 1d, e). Indeed, the beat-to-beat interval increased over the three beats following puff onset, and this increase was comparable across puff strengths (Supplementary Fig. 2e, f). Surgical removal of body parts and silencing experiments of external sensilla or chordotonal organs^55,56^ suggest that hair plate neurons and chordotonal organs are at least partially required for puff-induced cardiac deceleration (Supplementary Fig. 3). We also found that the air puff persistently increased the fraction of time walking, in line with previous reports^53,57^ (Fig. 1f, g; Supplementary Fig. 2c, d). A close inspection of the peri-stimulus dynamics revealed that the onsets of the cardiac and locomotor changes were approximately concurrent at puff onset across strengths (Fig. 1h). Furthermore, the peak of cardiac deceleration (roughly 0.2 – 0.5 s post puff onset) preceded that of locomotor increase (roughly 1 – 1.5 s post puff onset). Especially at lower puff strengths, locomotor response showed an additional transient dip, embedded in dominant and persistent locomotion, almost concurrently with (though, in the case of 50 mL/min, following) cardiac deceleration (Fig. 1h). Overall, cardiac and locomotor responses begin together, and the cardiac response reaches its peak before the locomotor response does. These data are consistent with the idea that cardiac and locomotor responses are coordinated, and that cardiac deceleration may causally contribute to the locomotor response. We therefore set out to identify the neurons responsible for cardiac deceleration and to test their causal role directly.

### II. Dopaminergic system is responsible for puff-induced cardiac deceleration

To aid in identifying neurons responsible for cardiac deceleration, we developed yet another paradigm to quantify the HR with higher throughput. In this paradigm, we drove the expression of GFP in the cardiomyocytes using the enhancer of the Hand gene^46^, to aid in heartbeat imaging (Fig. 2a, b). GFP signals in pericardiocytes (also known as nephrocytes) are likely due to their function to incorporate any molecules in the hemolymph^58^. The fly’s ventral side of the thorax and abdomen were glued to a slide glass, and the GFP signals in the conical chamber were acquired through the intact dorsal cuticle by confocal microscopy (Fig. 2a, b). The air puffs of 20 mL/min were applied frontally to the fly’s head through a glass pipette (Fig. 2a). Heartbeat was detected using our custom software (Supplementary Fig. 4a-c). A single trial consisted of 60 s of interval, 10 s of air puffs, and 60 s of recording, which was repeated twice for each fly with 90s interval (Fig. 2c). This experiment revealed that air puffs induce cardiac deceleration by roughly 10% (Fig. 2d-f; Supplementary Movie 2), consistent with our earlier finding (Fig. 1). Similar results were obtained when the air puff was applied for 1s (Supplementary Fig. 4d, e), 20s (Supplementary Fig. 4f, g), and in both sexes (Supplementary Fig. 4h, i).

**Fig. 2.**
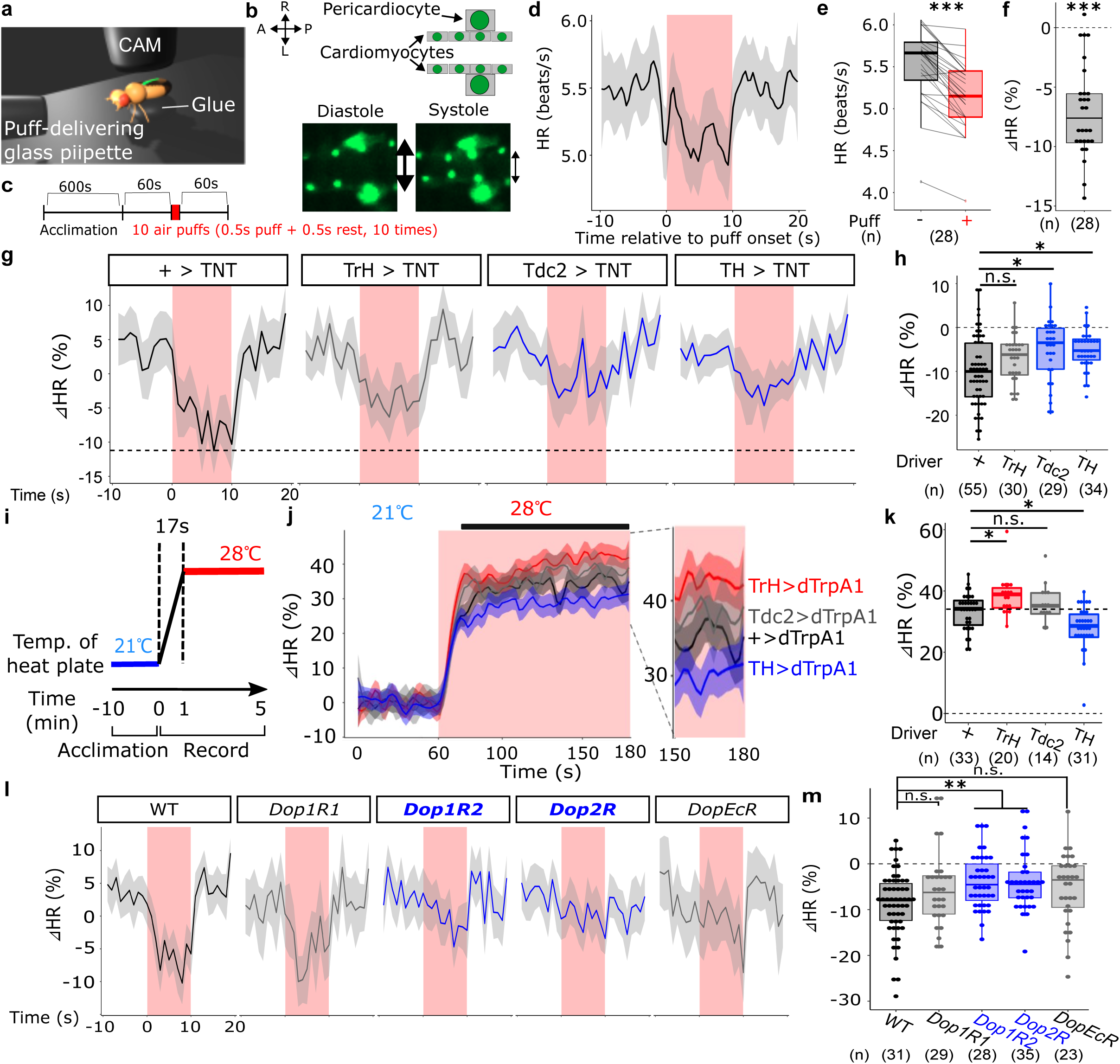
Dopamine system is responsible for puff-induced cardiac deceleration. **a.** Experimental setup. Hand enhancer drives the GFP expression in the cardiomyocytes (and in pericardiocytes). The fly was glued onto a slide glass, and the GFP signals were acquired through the intact cuticle. The air puffs were applied frontally to the fly’s head. **b.** Example heartbeat. (Upper) Cartoon of the acquired image. Two rows of small cardiomyocytes and one pair of large pericardiocytes are visible. (Lower) Diastole and systole can be distinguished by the changing distance between two rows of cardiomyocytes (arrows). **c.** The time course of the experiment. Air puffs of 0.5 s were applied at 1 Hz for 10 times (totalling 10 s). The trial was repeated twice for each fly, the results of which were averaged. **d.** Time course of the HR. Air puffs are applied during red-shaded window. Lines and shaded areas represent means and SEM, respectively. N = 28. **e.** The HR averaged during puff application window (“puff +”) vs 10 s before puff application (“puff -”). Each line corresponds to each fly. N = 28, ***p < 0.001, two-tailed *t*-test. **f.** Difference between puff+ HR and puff-HR, calculated for each fly. N = 28, ***p < 0.001, two-tailed *t*-test. **g.** Time course of the HR (% change from baseline defined as the average during the 10 s preceding puff onset). Air puffs were applied during the red-shaded time window. Lines and shaded areas represent means and SEM, respectively. TNT expression is driven by indicated drivers. “+” indicates driver-less control. Dashed line shows the lowest mean of “+” control during puffs. N = 55, 30, 29, 34, for +, TrH, Tdc2, TH, respectively. **h.** Change in HR, calculated for each fly as the % difference between the mean HR during the 10 s stimulation period and the mean HR during the 10 s baseline period preceding stimulation. N = 55, 30, 29, 34, for +, TrH, Tdc2, TH, respectively. *p < 0.05, n.s.: p > 0.05, Dunn’s test. **i.** Time course of the experiment. The heat plate beneath the slide glass warmed up the fly. The temperature of the slide glass is held at 21℃ for 10 min and ramped up to 28℃ for 5min. Note it took approximately 17 s for the temperature to ramp from 21℃ to 28℃. **j.** Time course of the HR, shown as % change from average HR at 21℃. Temperature was rampted from 21℃ to 28℃ during red-shaded window. **k.** % Changes in the HR (28℃ vs 21℃) from the baseline (average over initial 60 s) averaged over 75s -180s window (shown as black bar in j.). N = 33, 20, 14, 31, for +, TrH, Tdc2, TH, respectively. *p < 0.05, n.s.: p > 0.05, Dunn’s test. The upper dashed line shows the median of “+” control. **l.** Time course of the HR (% change from baseline defined as the average during the 10 s preceding puff onset). Air puffs were applied during the red-shaded time window. Lines and shaded areas represent means and SEM, respectively. N = 31, 29, 28, 35, 23, for +, WT, *Dop1R1*, *Dop1R2*, *Dop2R*, *DopEcR*, respectively. **m.** Change in HR, calculated for each fly as the % difference between the mean HR during the 10 s stimulation period and the HR during the 10 s baseline period preceding stimulation. N = 31, 29, 28, 35, 23, for +, WT, *Dop1R1*, *Dop1R2*, *Dop2R*, *DopEcR*, respectively. **p < 0.01, n.s.: p > 0.05, Dunn’s test.

Monoaminergic systems have repeatedly been implicated in HR regulation across species, including Drosophila^59–68^. We thus wondered whether monoaminergic systems might be involved in puff-induced cardiac deceleration. When we silenced with TNT the neurons expressing either serotonin, octopamine (insect equivalent of norepinephrine), and dopamine, we found that silencing of octopaminergic or dopaminergic neurons significantly suppressed puff-induced cardiac deceleration (Fig. 2g, h; Supplementary Fig. 5a, b). Note, however, that when we expressed in serotonergic neurons the inward-rectifying potassium channel Kir2.1, it did decrease the puff-induced cardiac deceleration (Supplementary Fig. 5c, d), suggesting that serotonergic neurons may also be indispensable for puff-induced cardiac deceleration. In line with this, a *loss-of-function* mutation of a serotonin synthetic enzyme *TrH* also showed decrease in the puff-induced cardiac deceleration (Supplementary Fig. 5e-g).

To further probe the roles of monoaminergic neurons, we next tested whether thermogenetic activation of these neurons can induce cardiac deceleration without air puffs. To this end, we expressed a heat-gated cation channel dTrpA1 in monoaminergic neurons, and shifted the temperature from 21℃ to 28℃ while monitoring the HR (Fig. 2i). This revealed that, while the HR of cold-blooded Drosophila increased as a function the temperature even in the driver-less control (Fig. 2j, black), such cardiac acceleration was suppressed by activation of dopaminergic neurons (Fig. 2j, blue) but not by octopaminergic neurons (Fig. 2j, gray). On the other hand, activation of serotonergic neurons exaggerated the heat-induced cardiac acceleration (Fig. 2j, red). The average HR during 28℃ period indeed revealed that the activation of dopaminergic neurons significantly decreased the HR compared to the driver-less control, while the activation of serotonergic neurons significantly increased the HR (Fig. 2k). Dopaminergic neurons were the only monoaminergic population whose silencing and activation consistently implicated them in driving cardiac deceleration (Fig. 2g-k). Serotonergic system, on the other hand, provided paradoxical results (Fig. 2g-k, Supplementary Fig. 5c-g): this could be because serotonergic system is required for threat-induced cardiac deceleration, while broad serotonergic activation outside this context possibly engages downstream circuits that promote cardiac acceleration. The role of dopaminergic neurons is further reinforced by *loss-of-function* experiments of dopamine receptors, which revealed that mutations of *Dop1R2* or *Dop2R* significantly decrease the puff-induced cardiac deceleration (Fig. 2l, m; Supplementary Fig. 5h, i). A mutation of *DopEcR* also showed a tendency to decrease the puff-induced cardiac deceleration albeit not quite reaching the statistical significance (Fig. 2l, m; Supplementary Fig. 5h, i).

Together, our data demonstrates that, while not excluding the contributions of serotonergic or octopaminergic systems, dopamine system is responsible for puff-induced cardiac deceleration.

### III. Two pairs of dopaminergic neurons in the brain, DA-WED neurons, are indispensable for cardiac deceleration

There are approximately 130 dopaminergic neurons scattered throughout the Drosophila brain^69,70^. To identify the subpopulation of dopaminergic neurons responsible for puff-induced cardiac deceleration, we turned to a host of dopamine-related GAL4 driver lines^69^ (Fig. 3a). Each of these drivers carries different bits of enhancers of dopamine-related genes, and presumably for this reason these drivers are known to label partially overlapping but distinct subpopulations of dopaminergic neurons^69^. When we drove the expression of TNT by these drivers, we found that neuronal silencing by 4 out of 10 drivers decreased puff-induced cardiac deceleration (R76F01-, R76F02-, R76F05-, R60F07-GAL4; Fig. 3b, blue plots) without significantly influencing the baseline HR (Supplementary Fig. 6a), suggesting that these drivers label the subset of dopaminergic neurons indispensable for puff-induced cardiac deceleration. Identification of these “hit” drivers allowed us to next employ a split-GAL4 approach to further narrow down the indispensable neurons (Fig. 3c). In this approach, GAL4 is divided into DBD and AD hemi-drivers, and functional GAL4 is reconstituted only in cells expressing both components ^71^ (Fig. 3c). When we tested all split combinations of the 4 “hit” enhancers, we found that only one combination, namely R76F05-DBD and R60F07-AD, significantly decreased the puff-induced cardiac deceleration (Fig. 3d), without significantly influencing the baseline HR (Supplementary Fig. 6b). Our immunohistochemical analysis revealed that, as reported by a previous report^69^, the split combination of R76F05-DBD and R60F07-AD labeled two different subpopulations of dopaminergic neurons: one subpopulation from the PPL1 cluster named PPL1-SMPγ, and another subpopulation from the PPM2 cluster named DA-WED (Fig. 3e), with additional one or two neurons occasionally labeled in some brains. We further sought to identify which of these two subpopulations is indispensable for puff-induced cardiac deceleration. Taking advantage of the recently-developed analysis pipeline that identify a list of candidate GAL4 drivers that may label any neuron of interest^72,73^, we obtained a list of candidate PPL1-SMPγ and DA-WED drivers. By testing several split combinations of enhancers used in these drivers, we could identify the split combinations that label one but not the other (Fig. 3f). We note that each of these combinations labels additional neurons, in either subesophageal zone (Fig. 3f, left) or optic lobes (Fig. 3f, right). Using these split drivers, we silenced either PPL1-SMPγ or DA-WED with TNT, and found that silencing of DA-WED significantly decreased puff-induced cardiac deceleration (Fig. 3g; Supplementary Fig. 6c). Together, our data suggests that DA-WED neurons are indispensable for puff-induced cardiac deceleration.

**Fig. 3.**
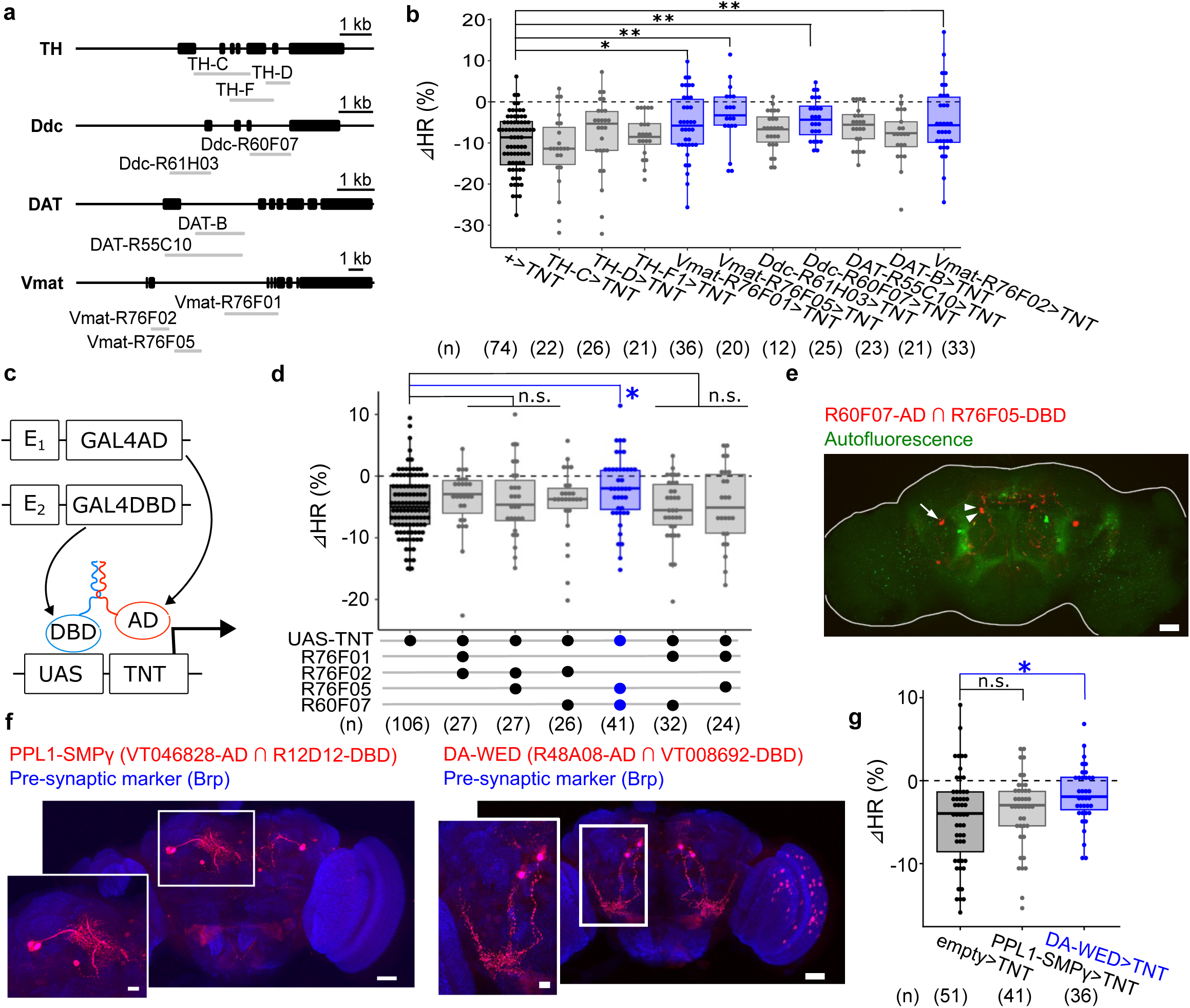
Two pairs of dopaminergic neurons in the brain are paritally indispensable for cardiac deceleration. **a.** AsSchematic of different DA-related GAL4 drivers. **b.** Change in HR, calculated for each fly as the % difference between the mean HR during the 10 s stimulation period and the HR HR during the 10 s baseline period preceding stimulation. N = 74, 22, 26, 21, 36, 20, 12, 25, 23, for +, TH-C, TH-D, TH-F1, Vmat-R76F01, Vmat-R76F05, Ddc-R61H03, Ddc-R60F07, DAT-R55C10, respectively. **p < 0.01, *p < 0.05, n.s. (data shown in gray): p > 0.05, Dunnett’s test. **c.** A schematic of split-GAL4 strategy. **d.** Change in HR, calculated for each fly as the % difference between the mean HR during the 10 s stimulation period and the HR HR during the 10 s baseline period preceding stimulation. N = 106, 27, 27, 26, 41, 32, 24, for driver-less control, R76F01 ∩ R76F02, R76F02 ∩ R76F05, R76F01 ∩ R76F02, R76F02 ∩ R76F07, R76F05 ∩ R60F07, R76F01 ∩ R60F07, R76F01 ∩ R76F05, respectively. *p < 0.05, n.s.: p > 0.05, Dunn’s test. **e.** Brain of R60F07-AD ∩ R76F01-DBD>mCherry (red). An arrow indicates the soma of PPL1-SMPγ, while arrowheads indicate the soma of DA-WED. Scale bar: 25 um. Similar results were obtained across 3 independent samples. **f.** Brains of VT046828-AD ∩ R12D12-DBD>mCherry and R48A08-AD ∩ VT008692-DBD>mCherry (red) immunostained with anti-neuropil marker nc82 (blue). Scale bars: 25 um (inlet: 5 um). Similar results were obtained across 5 independent samples for each genotype. **g.** Change in HR, calculated for each fly as the % difference between the mean HR during the 10 s stimulation period and the HR HR during the 10 s baseline period preceding stimulation. N = 51, 41, 36 for empty>TNT, PPL1-SMPγ>TNT, DA-WED>TNT, respectively. *p < 0.05, n.s.: p > 0.05, Dunnett’s test.

### IV. DA-WED neurons are responsible for puff-induced cardiac deceleration

While our data so far suggests the role of DA-WED neurons in cardiac deceleration, the split driver we used to label DA-WED also labeled neurons in the optic lobes (Fig. 3f; right). We thus further improved the labeling specificity through intersection strategy. In this strategy, the effector gene is expressed under the control of UAS sequence, but a “stop” sequence is inserted in between UAS and GFP, blocking the function of GAL4. Now, the expression of “molecular scissors” called flippase (FLP) is driven by a Tyrosine hydroxylase (TH) promoter (“TH-FLP”), which excises out the stop sequence. Hence, GAL4 can function only in cells in which TH promoter is active. This strategy, combined with our split GAL4 combination, indeed achieved near-perfect specificity for DA-WED neurons, apparently labeling no other cells throughout the body (Fig. 4a; Supplementary Fig. 6d, e). When we silenced DA-WED neurons with Kir2.1, it decreased puff-induced cardiac deceleration (Fig. 4b, c), without noticeably influencing the baseline HR (Supplementary Fig. 6f).

**Fig. 4.**
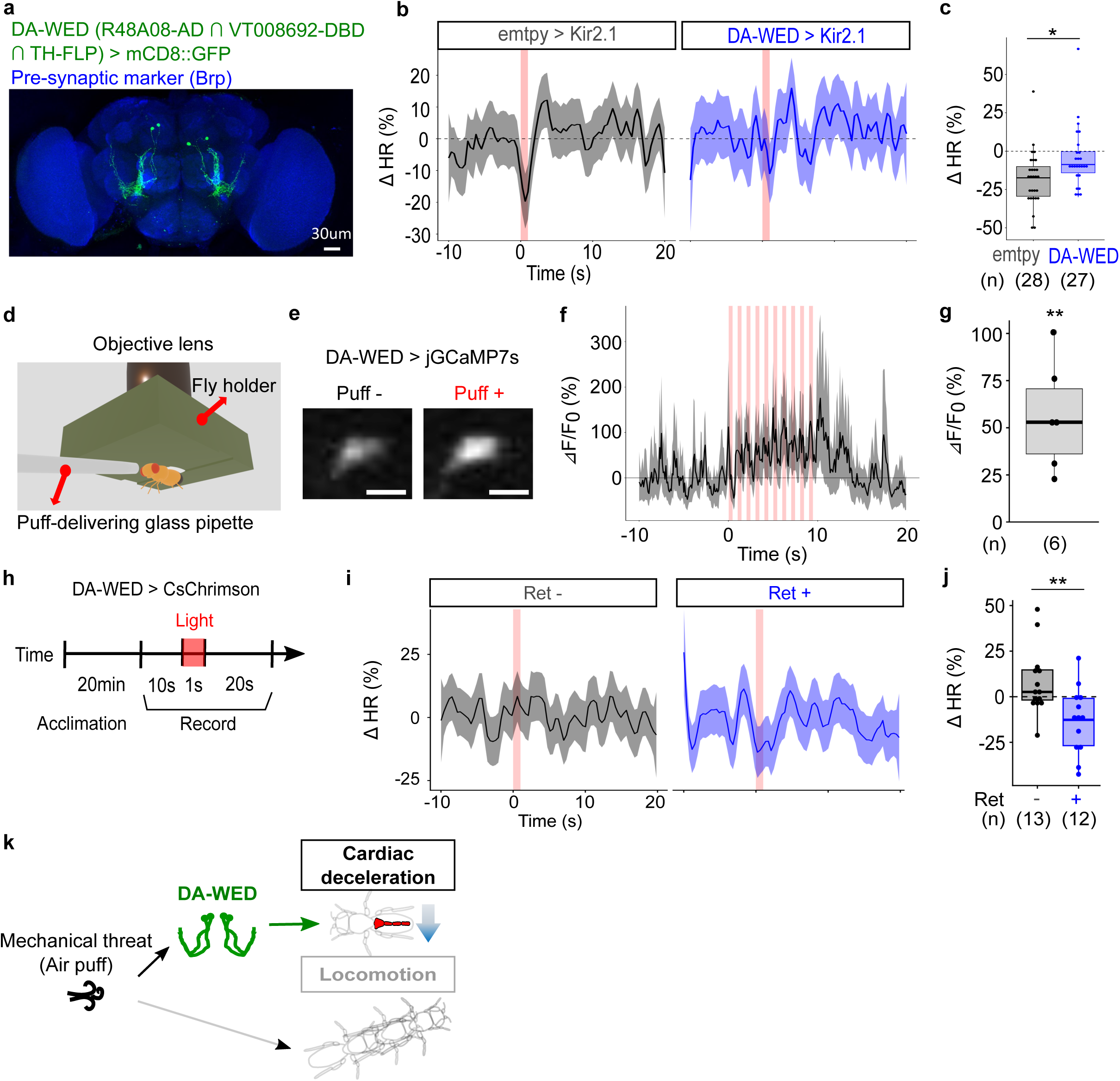
DA-WED neurons are responsible for puff-induced cardiac deceleration. **a.** The brain of R48A08-AD ∩ VT008692-DBD ∩ TH-FLP>mCD8::GFP (green) immunostained with anti-neuropil marker nc82 (blue). Scale bar: 30 um. Similar results were obtained across 5 independent samples. **b.** Time course of the HR (% change from baseline defined as the average during the 10 s preceding puff onset). Air puff was applied during the red-shaded time window. Lines and shaded areas represent means and SEM, respectively. N = 28, 27, for empty>Kir2.1, DA-WED>Kir2.1, respectively. **c.** Change in HR, calculated for each fly as the % difference between the mean HR during the 1 s following stimulation onset and the mean HR during the 10 s baseline period preceding stimulation. N = 28, 27, for empty>Kir2.1, DA-WED>Kir2.1, respectively. *p < 0.05, Two-tailed *t*-test. **d.** Experimental setup. **e.** Representative images of DA-WED neurons expressing GCaMP7s before and during puff application. Scale bar: 5μm. **f.** Time course of ΔF/F_0_. Puffs were applied at 1Hz for 10 times (over 10 s), as indicated by red-shaded boxes. N = 6 flies. **g.** ΔF/F_0_ averaged over 10s of puff application. When multiple neurons were recorded from the same fly (max 4 neurons per fly), results were averaged so that each dot represents one fly. N = 6 flies. two-tailed *t*-test. **p < 0.01. **h.** Time course of the experiment. After 20 min of acclimation, the heartbeat of the fly expressing light-gated cation channel CsChrimson in DA-WED neurons, with (“Ret +”) or without (“Ret -”) being fed with the co-factor all-trans retinal (ATR) for 2 days prior to experiment, was recorded for 10s, followed by 1s of light application, and 20s of recording. **i.** Time course of the HR (% change from baseline defined as the average during the 10 s preceding puff onset). Red light was shined during the red-shaded time window. Lines and shaded areas represent means and SEM, respectively. N = 19, 22, for Ret-, Ret+, respectively. **j.** Change in HR, calculated for each fly as the % difference between the mean HR during the 1 s following stimulation onset and the mean HR during the 10 s baseline period preceding stimulation. N = 13, 12, for Ret-, Ret+, respectively. *p < 0.05, two-tailed *t*-test. **k.** Model.

We next tested if DA-WED neurons function through dopamine. Our dopamine receptor mutant analysis (Fig. 2l, m) suggests that dopamine signaling through Dop1R2 and Dop2R is indispensable for cardiac deceleration. Consistent with this, we found that RNAi of a dopamine synthesis enzyme ple in DA-WED neurons reduced puff-induced cardiac deceleration (Supplementary Fig. 6g-i). It is thus plausible that DA released from DA-WED neurons binds to Dop1R2 and/or Dop2R to drive cardiac deceleration.

DA-WED neurons have previously been reported to drive protein preference behavior in protein-deprived flies^74,75^. Indeed, suppression of DA-WED neurons decreased protein preference (Supplementary Fig. 7a-d)^74,75^ in addition to cardiac deceleration (Supplementary Fig. 7e, Fig.4b, c). This prompted us to ask whether these neurons drive cardiac deceleration and protein preference through shared, or instead distinct, mechanisms. Interestingly, *Dop2R* mutant flies failed to show cardiac deceleration (Fig. 2l, m), but showed normal preference compared to the wildtype (Supplementary Fig. 7f, g). Similarly, suppression of FB-LAL neurons, through which DA-WED neurons drive protein preference^74,75^, failed to influence cardiac deceleration (Supplementary Fig. 7h, i). These results suggest that DA-WED neurons drive protein preference through FB-LAL neurons, while driving cardiac deceleration through Dop2R independent of FB-LAL. This hypothesis is further reinforced by our observations that protein deprivation, which drives protein preference (Supplementary Fig. 7j), fails to influence puff-induced cardiac deceleration (Supplementary Fig. 7k) nor the baseline HR (Supplementary Fig. 7l). Together, our data suggests that DA-WED neurons drive cardiac deceleration and protein preference through distinct mechanisms (Supplementary Fig. 7m).

We next wondered how DA-WED neurons might respond to air puffs. To this end, we expressed jGCaMP7s in DA-WED neurons, and imaged their calcium activities while applying a train of air puffs (Fig. 4d). We found that the calcium level of DA-WED neurons steadily increased as air puffs are applied (Fig. 4e, f; Supplementary Fig. 8a; Supplementary Movie 3). The increased jGCaMP7s signals returned to the baseline in less than 5 s when puffs ceased (Fig. 4f). Given the slow kinetics of jGCaMP7s^76^, the increased activity of DA-WED neurons likely returns to the baseline immediately when the puffs cease. While there are two pairs of DA-WED neurons in the brain, one pair being more medial than the other, we found no noticeable difference in medial vs lateral DA-WED neurons’ activities in response to puffs (Supplementary Fig. 8b-d). We note, though, that the medial neurons tended to show offset response-like activities more pronounced than their lateral counterparts (Supplementary Fig. 8c), potentially hinting at functional heterogeneity within DA-WED neurons. Overall, our data suggests that all four DA-WED neurons transiently get activated by air puffs.

To test if increased activities of DA-WED neurons can replace the air puff to induce cardiac deceleration, we photoactivated DA-WED neurons without puff application (Fig. 4h). For this, we expressed a light-gated cation channel CsChrimson in DA-WED neurons, and shined light on the fly’s head to activate these neurons. We found that the photoactivation of DA-WED neurons could induce cardiac deceleration by ∼10% in the retinal-fed test flies, but not in the retinal-unfed control flies (Fig. 4i, j; Supplementary Movie 3).

Together, our data establishes that DA-WED neurons are responsible for puff-induced cardiac deceleration (Fig. 4k).

### V. Cardiac interoception possibly contributes to threat-induced locomotion

Cardiac interoception has recently been implicated in modulating behavioral responses ^42–44^. We therefore asked whether cardiac deceleration is associated with changes in locomotion in *Drosophila*. To test this, we silenced DA-WED neurons using Kir2.1 to block puff-induced cardiac deceleration, and assessed locomotor output. We found that silencing DA-WED neurons markedly suppressed puff-induced locomotion compared to controls (Fig. 5a, b). Conversely, photoactivation of DA-WED neurons using CsChrimson promoted locomotion in the absence of puff (Fig. 5c, d). These findings suggest that DA-WED neurons coordinate cardiac deceleration with locomotor responses upon mechanical threat.

**Fig. 5.**
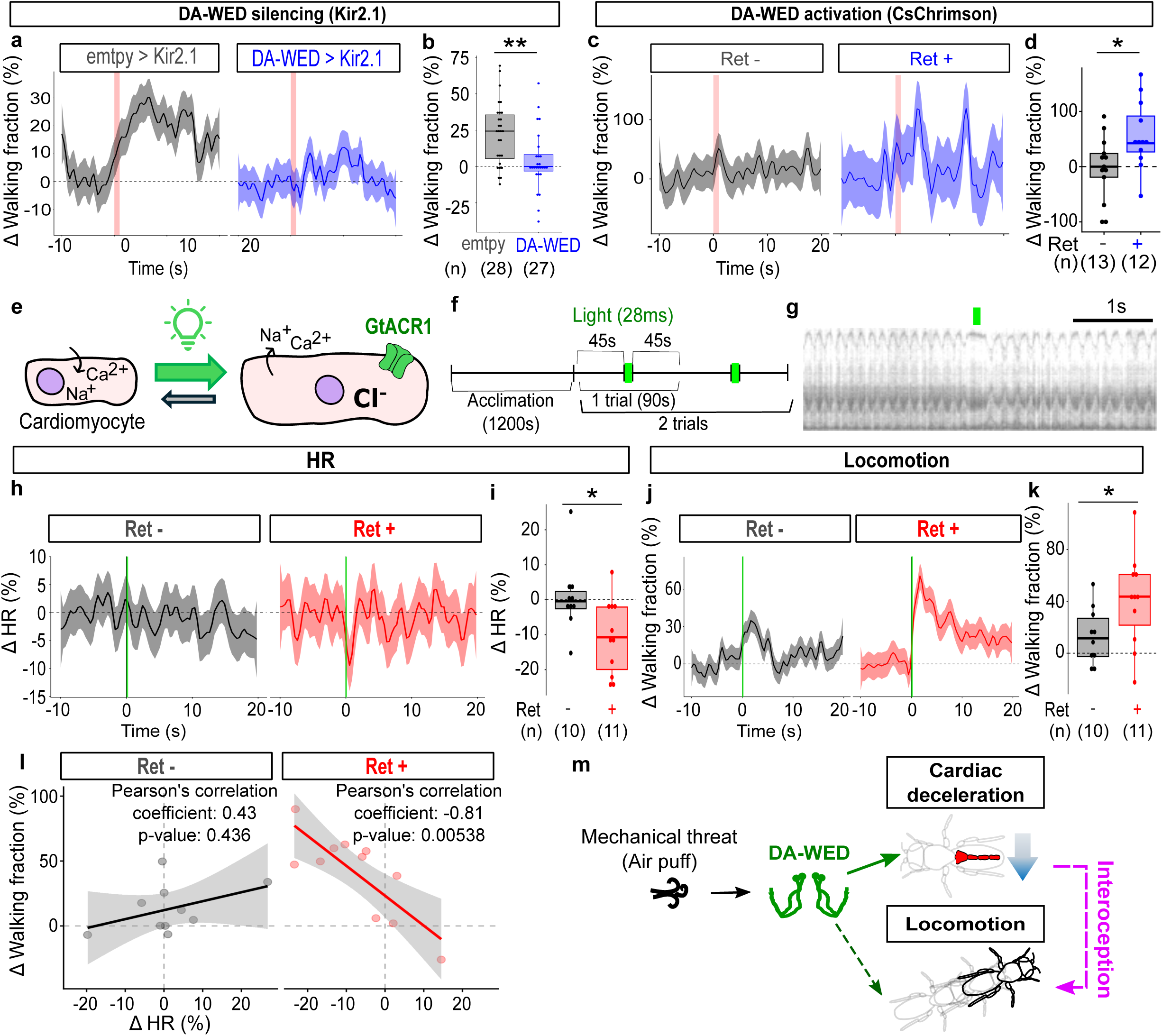
Cardiac interoception possibly contributes to threat-induced locomotion. **a.** Time course of the walking fraction (% change from baseline defined as the mean fraction during the 10 s preceding puff onset). An air puff was applied during the red-shaded time window. Lines and shaded areas represent means and SEM, respectively. N = 28, 27, for empty>Kir2.1, DA-WED>Kir2.1, respectively. **b.** Change in walking fraction, calculated for each fly as the % difference between the mean fraction during the 10 s following stimulation onset and the mean fraction during the 10 s baseline period preceding stimulation. N = 28, 27, for empty>Kir2.1, DA-WED>Kir2.1, respectively. **p < 0.01, Two-tailed *t*-test. **c.** Time course of the walking fraction (% change from baseline defined as the mean fraction during the 10 s preceding puff onset). Red light was applied during the red-shaded time window. Lines and shaded areas represent means and SEM, respectively. N = 13, 12, for ret -, +, respectively. **d.** Change in walking fraction, calculated for each fly as the % difference between the mean fraction during the 10 s following stimulation onset and the mean fraction during the 10 s baseline period preceding stimulation. N = 13, 12, for ret -, +, respectively. **p < 0.01, Two-tailed *t*-test. **e.** A schematic of the strategy. Cardiomyocytes beat by oscillating between shrunk (systole) and relaxed (diastole) states. Diastole is driven by efflux of cation and resultant hyperpolarization. We expressed a light-gated anion channel GtACR1 in cardiomyocytes using Hand-GAL4, thus artificially driving hyperpolarization by shining light. **f.** Time course of the experiment. After 20 min of acclimation, the heartbeat of the fly expressing GtACR1 in cardiomyocytes, with or without being fed with the co-factor all-trans retinal (ATR) for 2 days prior to experiment, was recorded for 90s, followed by a 28ms light pulse, and 45s of recording. The experiment was repeated twice for each fly. **g.** An example M-mode during optogenetic cardiac deceleration. Green bar indicates when the light was shined. **h.** Time course of the HR (% change from baseline defined as the average during the 10 s preceding puff onset). Green light was shined during the breen-shaded time window. Lines and shaded areas represent means and SEM, respectively. N = 10, 11, for ret -, ret +, respectively. **i.** Change in HR, calculated for each fly as the % difference between the mean HR during the 1 s following stimulation onset and the mean HR during the 10 s baseline period preceding stimulation. N = 10, 11, for ret -, ret +, respectively. *p < 0.05, two-tailed *t*-test. **j.** Time course of the walking fraction (% change from baseline defined as the mean fraction during the 10 s preceding puff onset). Light was shined during the green-shaded time window. Lines and shaded areas represent means and SEM, respectively. N = 10, 11, for ret -, ret +, respectively. **k.** Change in walking fraction, calculated for each fly as the % difference between the mean fraction during the 10 s following stimulation onset and the mean fraction during the 10 s baseline period preceding stimulation. N = 10, 11, for ret -, ret +, respectively. *p < 0.05, two-tailed *t*-test. **l.** Correlations between changes in walking fraction and HR. N = 10, 11, for ret -, ret +, respectively. **m.** Model. DA-WED neurons promote locomotion, through and/or bypassing interoception of the heartbeat (dashed arrows).

An alternative interpretation is that DA-WED neurons function as mere puff detectors that broadly regulate puff-evoked behaviors. If this were the case, manipulating these neurons should affect all behavioral outputs induced by puff. In addition to promoting locomotion, air puff suppressed grooming predominantly within 3 s of puff onset, consistently across puff strengths; grooming during the later 10–20 s window was variable, either remaining suppressed, returning to baseline, or reversing to an increase depending on puff strength (Supplementary Fig. 9a–c). Silencing DA-WED neurons failed to disrupt puff-induced grooming suppression (Supplementary Fig. 9d–f), despite markedly reducing cardiac deceleration (Fig. 4b, c) and locomotion (Fig. 5a, b). Likewise, photoactivation of DA-WED neurons failed to suppress grooming (Supplementary Fig. 9g–i), while robustly inducing cardiac deceleration (Fig. 4i, j) and locomotion (Fig. 5c, d). Thus, manipulation of DA-WED neurons does not reproduce the full behavioral repertoire triggered by puff. These findings argue against the idea that DA-WED neurons function as mere puff detectors and instead support a more specific role in coordinating cardiac and locomotor responses.

To directly test whether cardiac deceleration can influence locomotion, we optogenetically induced cardiac deceleration of a magnitude comparable to the puff-induced response by expressing the light-gated anion channel GtACR1 in cardiomyocytes using Hand-GAL4 (Fig. 5e). We selected an anion channel to prolong diastole, unlike a previous report using a cation channel^77^, as puff-induced cardiac deceleration is characterized by elongated diastolic intervals (Fig. 1c). To approximate the 20–23 ms beat-to-beat prolongation evoked by mechanical threat (Fig. 1; Supplementary Fig. 2e, f), we used 28-ms (2-frame) green light pulses. The light pulses transiently arrested the heart in diastole (Fig. 5f, g; Supplementary Movie 5), reducing HR by ∼10% (Fig. 5h, i), comparable to puff-and DA-WED-induced deceleration (Fig. 1d, e; Fig. 4i, j). Notably, optogenetic induction of cardiac deceleration in cardiomyocytes—independent of central dopaminergic activation—was accompanied by increased locomotion (Fig. 5j, k). In contrast, this manipulation failed to alter grooming behavior (Supplementary Fig. 9j-l), suggesting a selective cardiac influence on locomotion. Furthermore, optogenetically induced changes in HR and locomotion were strongly anti-correlated fly-wise in the retinal-fed test group (Pearson’s correlation coefficient: -0.81, p-value: 0.00538) but not in the control group without retinal (Pearson’s correlation coefficient: 0.43, p-value: 0.436) (Fig. 5l). This contrasts with puff-induced responses, in which changes in HR and locomotion were not significantly correlated (Supplementary Fig. 10a). This lack of correlation may reflect contributions from cardiac-state-independent locomotor pathways. These observations are consistent with the possibility that cardiac deceleration may influence locomotor responses upon mechanical threat (Fig. 5m; Supplementary Fig. 9m). One limitation is that our optogenetically induced cardiac pattern (Fig. 5g; Supplementary Fig. 10b, c) is not necessarily identical to its mechanically induced counterpart (Fig. 1c; Supplementary Fig. 10d, e), which could have influenced locomotion. We also note that retinal-unfed controls showed a modest locomotor increase following green-light illumination (Fig. 5j, k), leaving open whether cardiac deceleration directly promotes locomotion or potentiates a light-evoked locomotion. In either case, our data support a modulatory influence of cardiac state on locomotor output.

Altogether, these results identify a group of dopaminergic neurons that mediate threat-induced cardiac deceleration and point to a potential role of cardiac dynamics in shaping locomotor responses during threat.

## DISCUSSION

Here we demonstrate that mechanical threat engages dopaminergic neurons that induce rapid cardiac deceleration and is associated with increased locomotion. Air puff stimulation elicited both cardiac deceleration and locomotor activation (Fig. 1), and we identified two pairs of dopaminergic neurons, termed DA-WED neurons, as key mediators of the cardiac response (Fig. 4). Silencing these neurons attenuated threat-induced cardiac deceleration and locomotion, whereas optogenetic activation of DA-WED neurons elicited both responses in the absence of threat (Fig. 4, Fig. 5a-d). Moreover, direct optogenetic cardiac deceleration was accompanied by increased locomotion (Fig. 5e-l), suggesting that cardiac deceleration may influence behavioral responses. Together, these findings support a model in which descending dopaminergic control of cardiac activity contributes to the coordination of physiological and behavioral responses to threat, potentially via interoceptive feedback from the heart.

### Cardiac deceleration is coordinated with defensive locomotion

Our data show that air puff stimulation induces cardiac deceleration together with increased locomotion (Fig. 1). Across puff strengths, locomotion scaled with stimulus strength, whereas the cardiac deceleration did not: measured as time-binned HR, it was less pronounced at stronger puffs, while measured as the beat-to-beat interval it was comparable across strengths (Fig. 1d, e; Supplementary Fig. 2e, f). This suggests that cardiac deceleration is likely not simply a byproduct of locomotion.

Although cardiac deceleration is often associated with freezing in mammals^1^, cardiac deceleration has also been observed during early threat detection and orienting responses^78^, suggesting that it reflects engagement of a defensive processing rather than motor inhibition *per se*. Consistent with this view, dissociation between threat-evoked cardiac modulation and freezing has been reported across taxa. In mammals, for example, disruption of autonomic output can abolish cardiac deceleration without eliminating freezing^79^. Similarly, in decapod crustaceans and *Drosophila*, visual stimuli elicit reproducible and time-locked changes in HR that accompany both freezing and fleeing^8,9,80^. While these studies establish that cardiac modulation can be dissociated from freezing, our findings additionally demonstrate that cardiac deceleration can co-occur with a predominantly locomotor response to mechanical threat and can itself causally contribute to locomotion (Fig. 5e-m). Cardiac deceleration is therefore not restricted to passive defensive states, but can act upstream of active locomotor responses.

### Dopaminergic neurons in the brain are responsible for puff-induced cardiac deceleration

The present study identifies two pairs of dopaminergic neurons in the brain, termed DA-WED neurons, as key mediators of puff-induced cardiac deceleration. Silencing/ photoactivating DA-WED neurons disrupts/ mimics puff-induced cardiac deceleration, respectively (Fig. 4b, c, h-j). Furthermore, DA-WED neurons exhibit calcium elevation upon air puffs (Fig. 4d-g). Moreover, knockdown of the dopamine synthetic enzyme *ple* in DA-WED neurons suppresses puff-induced cardiac deceleration (Supplementary Fig. 6g–i), suggesting that dopamine release from these neurons is required for cardiac deceleration.

Circulatory dopamine has long been associated with cardiac acceleration across species^81^. In *Drosophila*, pharmacological application of DA to the exposed heart accelerates the heartbeat^60–63^, seemingly at odds with our findings. However, the responsible source of circulatory DA or DA receptor, as well as whether DA directly acts on cardiomyocytes or instead through some yet unidentified cells, remain elusive. In mammals, cardiomyocytes *per se* synthesize DA and express DA receptors, hinting at autocrine mechanism^81^. Circulatory DA also increases the levels of epinephrine and norepinephrine, complicating the interpretation of “DA-induced” cardiac acceleration^82^. From our ple RNAi experiment (Supplementary Fig. 6g-i), DA released from DA-WED neurons appears indispensable for puff-induced cardiac deceleration. Notably, neurites of DA-WED neurons remain confined to the brain, suggesting that DA-WED neurons engage circulation or yet unidentified neuronal circuitry to indirectly modulate the heartbeat. Circulation is unlikely to mediate DA-WED neurons’ function, however, given that the air puff and DA-WED activation both induce cardiac deceleration within a second (Fig. 1, Fig. 4). Interestingly, DA acting as a local neuromodulator to drive cardiac deceleration has been documented in mammals by a few literatures, either in the ventral tegmental area (VTA) or lateral habenula (LHb)^83,84^. Together, our findings and previous literature suggest that dopamine may act not only as a circulating hormone that accelerates HR, but as a centrally deployed neuromodulator capable of driving cardiac deceleration during defensive states.

### DA-WED neurons coordinate cardiac and behavioral threat responses

Across species, threatening stimulation elicits cardiac and behavioral responses in an orchestrated fashion. Cardiac interoception is postulated to serve as key feedback to link cardiac and behavioral responses^40^. Cardiac interoception is mediated by sensory neurons innervating the heart and aortic arch^40^, which convey information to ACC, insular, and prefrontal cortex^41^ through brain stem NTS and hypothalamic PVN^41^. These ascending pathways have been proposed to modulate learned fear^42^, anxiety-like behavior^43^ or sleep^44^. Although models propose that descending control of the heart and ascending interoceptive feedback jointly shape defensive behavior, experimental validation of this bidirectional framework within a single system has been limited.

Our results provide partial support for such a model. We show that (1) mechanical threat induces cardiac deceleration and locomotion (Fig. 1), (2) DA-WED neurons in the brain are responsible for threat-induced cardiac deceleration and locomotion (Fig. 4; Fig. 5a-d), (3) optogenetic cardiac deceleration is accompanied by increased locomotion (Fig. 5e-l). Our data points to a scenario in which DA-WED neurons induce cardiac deceleration upon mechanical threat, which may in turn influence threat-induced locomotion (Fig. 5m). We note, however, that our data does not exclude the possibility that DA-WED neurons might influence locomotion at least partially independent of its role in cardiac deceleration.

Notably, the behavioral consequences of cardiac deceleration appear to be selective rather than global. Although air puff stimulation suppressed grooming, neither silencing nor activation of DA-WED neurons significantly affected grooming responses (Supplementary Fig. 9a-i), despite robust effects on cardiac deceleration and locomotion (Fig. 4, Fig. 5). Similarly, direct optogenetic induction of cardiac deceleration failed to influence grooming behavior (Supplementary Fig. 9j-l). These dissociations indicate that cardiac feedback does not uniformly modulate defensive behaviors but preferentially facilitates locomotor output. Such selectivity is consistent with the possibility that interoceptive signals from the heart may bias specific behavioral strategies rather than simply increasing overall arousal.

### Cardiac dynamics can modulate behavior

Our findings echo a growing body of work linking cardiac dynamics to behavior across distant taxa. In larval zebrafish, cardiac dynamics and behavior are coupled during threat^85^, and dedicated vagal sensory neurons encode HR, including its decreases, through a stimulus-invariant population code^86^. In mice, optogenetically induced tachycardia can drive anxiety-like behavior^87^. And here we find that cardiac deceleration accompanies, and can promote, locomotion in *Drosophila*. Cardiac influence on behavior thus spans insects, fish, and mammals.

How cardiac state might be sensed in *Drosophila* is unknown, and several routes are conceivable. Drosophila heart receives rich neuronal innervation^51,52^, although whether these neurons relay sensory information from the heart is unknown. If so, neurons could directly detect the heartbeat, analogous to the cardiac afferents that mediate cardiac interoception in vertebrates^86,87^. Alternatively, the heartbeat could be sensed mechanically or through gas-exchange-related cues. In mammals, the cardiac pressure pulse can be detected directly within the brain by mechanosensitive central neurons^88^. In zebrafish, cardiac activity can causally influence behavior through its oxygen supply^85^. In insects, although the circulatory architecture differs, the heartbeat contributes to pressure changes in the hemocoel and tracheal system and resultant gas exchange^89,90^, which could in principle be detected. While present study does not distinguish among these possibilities, establishing how cardiac state is sensed would provide a foundation for dissecting how central commands and peripheral physiological states interact to coordinate survival behaviors.

### Limitations of this study

While our data are consistent with a model in which DA-WED neurons couple cardiac deceleration to locomotor responses upon mechanical threat, several questions remain. Although cardiac deceleration can promote locomotion, we have not directly tested whether it is required for DA-WED–evoked locomotion, leaving open the possibility of parallel downstream pathways. In addition, the sensory mechanisms that relay cardiac state to locomotor circuits in *Drosophila* remain unidentified. Finally, despite the rapid timescale and behavioral specificity of our manipulations, we cannot fully exclude indirect physiological contributions secondary to altered cardiac dynamics. Future dissection of descending and ascending circuits will clarify these mechanisms.

Overall, we identify a discrete population of dopaminergic neurons that drive rapid cardiac deceleration in response to mechanical threat. We further show that these neurons coordinate cardiac responses and are associated with behavioral components of the defensive response, potentially by engaging interoceptive feedback from the heart. Our findings establish *Drosophila* as a tractable model for dissecting how exteroceptive and interoceptive signals are integrated to couple organ physiology with adaptive behavior, opening new avenues for understanding the neural logic by which internal and external states are coordinated to promote survival.

## ONLINE METHODS

### Fly stocks

The following strains were obtained from Bloomington Stock Center (Indiana University): TrH-GAL4 (#38388), Tdc2-GAL4 (#9313), TH-GAL4 (#8848), UAS-TNT (#28838), UAS-Kir2.1::EGFP (#6596), UAS-CsChrimson::mVenus (#55135), UAS-dTrpA1 (#26263), UAS-mCD8::GFP (#32186), UAS-RedStinger (#8547), UAS-IVS-jGCaMP7s (#79032), UAS-GFP RNAi (#41560), UAS-Ple RNAi (#65875), VT046828-AD (#73115), R12D12-DBD (#69213), R48A08-AD (#70707), VT008692-DBD (#86909), empty-AD; empty-DBD (#79603), R70G12-GAL4 (#39552), R38B08-AD (#71032), R81E10-DBD (#68529),VT043775-DBD (#73728), R74C07-GAL4 (#39847), Aes1N-GAL4 (#39847), iav-GAL4 (#52273), R48A07-GAL4 (#50340), UAS-mCherry (#52267), Hand-GAL4 (#48396).

The following strains were obtained from Kyoto Drosophila Stock Center (Kyoto Institute of Technology): Tes1N-GAL4 (#103660).

Hand-GFP was a gift from Dr. Zhe Han (The University of Maryland).

UAS-FRT-stop-FRT-Kir2.1, and UAS-FRT-stop-FRT-CsChrimson::mVenus were gifts from Dr. Barry Dickson (The University of Queensland).

TH-C-GAL4, TH-D-GAL4, TH-F1-GAL4, Vmat-R76F01-GAL4, Vmat-R76F05-GAL4, Ddc-R61H03-GAL4, Ddc-R60F07-GAL4, DAT-R55C10-GAL4, Vmat-R76F01-AD, Vmat-R76F05-AD, Ddc-R60F07-AD, Vmat-R76F01-DBD, Vmat-R76F05-DBD, Ddc-R60F07-DBD, TH-FLP, UAS-FsF-mCD8GFP were gifts from Dr. Mark Wu (Johns Hopkins University).

*Dop1R1-attP*, *Dop1R2-attP*, *Dop2R-attP*, *DopEcR-attP, TrH-attP* were gifts from Dr. Yi Rao (Peking University).

UAS-GtACR1::tdTomato (III) was gift from Dr. Tetsuya Tabata (The University of Tokyo).

Genotypes and sample sizes used in this study are summarized in Supplementary Data 1.

### Fly husbandry

Flies were maintained on conventional cornmeal-agar-molasses medium under a 9AM:9PM light/ dark cycle at 25 °C and 60 +/-5 % humidity. Flies were collected 0–1 day post eclosion and housed in a group of 9-10 for 2-5 days before testing unless otherwise stated. Male flies were used throughout except for Supplementary Fig. 4h, i, in which both male and female were used. All experiments were carried out between 10 am and 8 pm at 25 +/-0.8 C and 60 +/-5 % humidity.

### Heartbeat and behavior recording

Flies were tethered above an air-supported foam ball (8 mm diameter; Yuzawaya, Japan) mounted in a custom sphere holder. Airflow was continuously monitored using a digital flowmeter (MF-FP10N-H06-010, Horiba Estec Inc.) and maintained at 500 ± 10 mL/min. Behavioral imaging was performed from the right side under 850 nm infrared illumination using a CMOS camera (GE60, Himawari; 640 × 480 pixels). Simultaneously, cardiac activity was imaged from the dorsal side through the intact cuticle under 850 nm illumination using a second CMOS camera (CS135MU, Thorlabs; 800 × 500 pixels). Behavioral and cardiac imaging streams were externally triggered and synchronized at 70 Hz using an Arduino Uno board.

Immediately prior to the experiment, flies were briefly (< 1 min) cold-anesthetized, and the dorsal thorax and abdomen were glued to a coverslip using UV-curable adhesive (Bondic BD-SKCJ) polymerized with 405 nm UV light (OSV5XME1C1E, Akizuki Electronics Inc.). The coverslip was positioned above the air-supported ball, allowing the fly to freely walk on the ball. Each recording session, except for DA-WED optogenetic activation experiment, consisted of a 20 min acclimation period followed by puff-response testing (950 s). During puff-response testing, a 500 ms air puff was delivered once every 90 s for a total of 10 puffs. Recording continued for 30 s after the final puff. DA-WED optogenetic activation experiment consisted of a 20 min acclimation period followed by photoactivation-response testing (950 s), in which a 1 s 617nm light was delivered once every 90 s for a total of two (See “Optogenetic activation of DA-WED neurons” below), to avoid potential desensitization of CsChrimson.

### Heartbeat (without behavior) recording

Cardiomyocytes expressing GFP were imaged dorsally through the intact cuticle. Images (104 × 50 pixels) were acquired at 24 Hz using a Leica SP8 confocal microscope equipped with a 10× objective lens (NA 0.4).

Immediately before recording, flies were briefly (< 1 min) cold-anesthetized, and the ventral thorax and abdomen were glued to a glass slide using UV-curable adhesive (Bondic BD-SKCJ) polymerized with 405 nm UV light (OSV5XME1C1E, Akizuki Electronics Inc.). The slide was positioned beneath the objective lens for imaging. Each recording session consisted of a 10 min acclimation period followed by puff-response testing (350 s total). A single puff-response test consisted of 1 min baseline recording, followed by a train of 10 air puffs delivered at 1 Hz (10 s total), and a subsequent 1 min post-stimulation recording. For each fly, the puff-response test was repeated twice with a 90 s inter-trial interval.

### Automatic heartbeat detection

To automatically detect individual heartbeats, we developed custom software tools. For infrared (IR)–based heart imaging obtained during simultaneous recordings of cardiac activity and behavior, the detection pipeline (Supplementary Fig. 1a) computed optical flow for each pixel between consecutive video frames. These optical flow measurements were used to quantify global cardiac relaxation (diastole) and contraction (systole). Time points corresponding to maximal contraction were identified as systolic peaks, and peak counts were used to compute heart rate (HR). The precision of the automated detection was evaluated by comparing both absolute HR values and percent changes in HR during air-puff stimulation relative to baseline with manual annotations. These analyses demonstrated that the automated method detected both absolute HR and stimulus-evoked HR changes with accuracy indistinguishable from manual scoring (Supplementary Fig. 1b, c).

To detect heartbeats in confocal microscopy images of GFP-expressing cardiomyocytes (recorded in the absence of behavioral measurements; Supplementary Fig. 4a), each frame was subtracted from the subsequent frame to compute pixel-wise differences between consecutive image pairs. This procedure generated positive and negative pixel values in regions surrounding the cardiomyocytes during cardiac contraction (systole) and relaxation (diastole). During systole, pixels within the heart tube exhibited positive values corresponding to the current positions of cardiomyocytes, whereas pixels outside the heart tube exhibited negative values corresponding to their previous positions; the opposite pattern was observed during diastole. Systolic and diastolic phases were distinguished by: (i) calculating the standard deviation of the positions of positive-value pixels along the axis perpendicular to the heart tube (the y-axis in Supplementary Fig. 4a), denoted SD(t); (ii) calculating the standard deviation of the positions of negative-value pixels along the same axis, denoted SD(t–1); and (iii) computing the difference SD(t) − SD(t–1). Negative values of SD(t) − SD(t–1) indicated systole, whereas positive values indicated diastole (Supplementary Fig. 4a). Time points corresponding to maximal contraction were identified as systolic peaks, and peak counts were used to compute HR. The precision of this algorithm was evaluated as described above. Although absolute HR values were slightly overestimated (Supplementary Fig. 4b), stimulus-evoked changes in HR were captured with accuracy indistinguishable from manual annotation (Supplementary Fig. 4c).

To obtain a continuous HR time course, peak counts were binned into 0.2 s intervals, and the number of beats per bin was calculated for each fly and trial. For visualization of HR dynamics relative to stimulus onset, HR traces were aligned to puff onset (time 0) and averaged across repeated trials within each fly. In experiments with a single air puff, puff-induced HR modulation was quantified by comparing the mean HR during the 1 s following puff onset with the mean HR during the 10 s baseline period preceding puff onset. Percent change was calculated as [(HR_post − HR_baseline) / HR_baseline] × 100. In experiments involving 10 puffs delivered over a 10 s window, HR modulation was quantified as the mean HR during the entire 10 s stimulation period relative to the 10 s pre-stimulus baseline.

### Automatic behavioral identification

To automatically identify behavior on a frame-by-frame basis, videos were analyzed offline using a custom Python-based pipeline. Body-part coordinates—including the joints of each leg, the eye, neck, haltere, and the rostral tip of the abdomen on the right side of the fly—were tracked using SLEAP software^54^. The tracked coordinates were subsequently transformed to normalize body length and orientation across individuals. Using these standardized coordinates, the speed of each body part was calculated for each pair of consecutive frames. Both the standardized coordinates and corresponding speed measurements were then provided as inputs to a custom support vector machine (SVM)–based classifier, which labeled each frame as “groom,” “walk,” or “halt.” The SVM was trained using the standardized body-part coordinates and speed features as predictors, with manually annotated behavioral labels serving as the dependent variables. For each fly and each time point relative to puff onset, we calculated the fraction of frames classified as walking or grooming across 10 repeated trials. For visualization of behavioral dynamics, these fraction time courses were aligned to puff onset (time 0) and averaged across trials within each fly. Puff-induced changes in walking and grooming were quantified by comparing the mean fraction during the 10 s and 3 s, respectively, following puff onset with the mean fraction during the 10 s baseline period preceding puff onset. Percent change was calculated as [(fraction_post −fraction_baseline) / fraction_baseline] × 100.

### Mechanical stimulation

Mechanical stimulation was delivered through a glass pipette (tip diameter of ∼2 mm), placed ∼2mm in front of the fly, connected with air hose to a compressor (EARTH MAN, Takagi, Japan) via a solenoid valve (EXA-C6-02C-4, CKD). The solenoid valve was controlled by an Arduino Uno board, which in turn received connection from a desktop computer running the python program which orchestrated the video recordings and puff application. The mechanical stimulation was generated by alternating the solenoid valve between 500 ms-open and 500 ms-closed states (1 Hz). The mechanical stimulation lasted for either 1s, 10s, or 20s, as specified in each figure. The air flow was monitored with a digital flowmeter (MF-FP10N-H06-010, Horiba estec. Inc) and was adjusted to either 20 mL, 50 mL, or 200 mL/min, as specified in each figure.

### Optogenetic activation of DA-WED neurons

Flies expressing CsChrimson, a red-shifted channelrhodopsin variant, in DA-WED neurons were raised on food containing 500 µM all-trans retinal for 2 -3 days, in the dark, prior to testing. Photoactivation was performed by delivering a 1s LED light pulse (617nm, Thorlabs) to the fly’s head. The output for LED was measured with a power meter (S120C and PM120D, Thorlabs) at a position corresponding to the fly’s head, and was adjusted to 1 μW/mm^2^. Light stimuli were triggered by python through an Arduino board connected to the LED driver.

### Optogenetic cardiac deceleration

Flies expressing GtACR1, a light-gated anion channel, in cardiomyocytes were raised on food containing 500 µM all-trans retinal for 2 -3 days, in the dark, prior to testing. Photoactivation was performed by delivering a 28ms-long LED light pulse (530nm, Thorlabs) to the fly’s heart through the objective lens. Because mechanical threat transiently elongated beat-to-beat intervals typically by 20 -23ms, we chose 28-ms (2-frame, at our 70-Hz acquisition rate) stimulation to best mimic this effect, over 14 (1-frame) or 42 (3-frame) ms (Fig. 1; Supplementary Fig. 2b, e, f).The light intensity was adjusted, for each fly prior to the recording, to the minimum intensity required for a light pulse of roughly 2s to briefly stop the heartbeat. After the recording, the output for LED was measured with a power meter (S120C and PM120D, Thorlabs) at a position corresponding to the fly’s heart, and ranged from 9.88 – 21.8μW/mm^2^. Light stimuli were triggered by python through an Arduino board connected to the LED driver.

### Immunohistochemistry

The following antibodies were used: mouse nc82 (1:10 Developmental Studies Hybridoma Bank), goat anti-mouse Alexa 633 (1:100, Molecular Probe #A21050). Whole brain immunohistochemistry was performed as described previously^91,92^. Briefly, brains were dissected out in 0.3 % PBST, and were fixed in 2 % paraformaldehyde/ PBS for 90 min at room temperature. Brains were then washed in 0.3 % PBST, for > 3 h, and were blocked in 10 % normal goat serum/ 0.3 % PBST for 30 min. Brains were incubated in the primary antibodies in 10% normal goat serum/ 0.3% PBST at 4 °C for 2 days. Brains were then washed in 0.3 % PBST for > 5 h, and were blocked in 10 % normal goat serum/ 0.3 % PBST for 30 min. Brains were then incubated in the secondary antibodies in 10 % normal goat serum/ 0.3 % PBST at 4℃ for 2 days. Afterward, brains were washed in 0.3 % PBST for > 5 h, and incubated in 50 % glycerol/ 0.3 % PBST for 2 h at 4 °C. Lastly, brains were mounted on slide glass in Vectashield (Vector Laboratories). Images were taken with Leica TCS SP8 confocal microscope, and processed in ImageJ (NIH) software.

### Calcium imaging

Two-photon calcium imaging was conducted as described previously^53^ with a few modifications. Two-photon microscopy (Bergamo II, Thorlabs) with a 16x objective lens (16x CFI LWP Plan Flour Objective, Nikon) and with near-infrared excitation (930 nm, Mai Tai 2, SpectraPhysics. Inc., Mountain View, CA) was used. Cuticle of the top of the flies’ head and the fat and trachea beneath it were surgically removed. The exposed brain was submerged in saline solution. The extracellular saline had the following composition (in mM): 103 NaCl, 3 KCl, 5 HEPES, 10 trehalose, 10 glucose, 7 sucrose, 26 NaHCO_3_, 1 NaH_2_PO_4_, 1.5 CaCl_2_, 4 MgCl_2_. We identified DA-WED neurons using the baseline fluorescence of GCaMP. Images were acquired from 1 plane at 152 frames per s (160 x 96 pixels, pixel size = 0.388 µm, dwell time = 0.246 µs). During calcium imaging, a large fraction of the fly’s body was glued to the holding plate, and part of the fly’s eye compounds was covered with the holder, and the fly was hung in the air. Each fly underwent the sequence of stimuli per fly once or twice, to scan DA-WED neurons on both hemispheres whenever possible. The time courses of ΔF/F_0_ were calculated by, for each bin, taking the percent difference from F 10s prior to the onset of the initial puff application (t = -10 through 0s in Fig. 4f).

### Surgery

In search of sensory organs indispensable for air puffs to induce cardiac deceleration, halteres, wings, legs, and a3 segments of the antennae were surgically removed using a pair of forceps on ice just prior to testing. Flies that underwent cold anesthesia just prior to testing served as controls.

### Feeding preference test

Two days post eclosion male flies were protein deprived by maintaining on 200 mM sucrose (vials with a kimwipe soaked with 2mL of sucrose solution) for 8 days (kimwipe is exchanged every other day). We prepared 7.5% yeast and 200mM sucrose in 0.9% of agar, color-labeled with either blue (brilliant blue FCF) or red (sulforhodamine B) tasteless food dyes. The two types of food mix, 1mL each, were then aliquoted into 35mm petri dish separated by a 5×5×35mm acrylic bar into two chambers. A group of 70 flies starved for 3 hours in an empty vial with a kimwipe soaked with 2 mL of water were cold-anesthetized and transferred into petri dish at ZT2 and allowed to feed for 2 hours in the dark. The flies were then killed by transferring the dish to -30C, upside-down, and scored by examining the color of their abdomen. Preference Index (PI) for yeast was calculated as (# flies consuming yeast -# flies consuming sucrose) / (total # flies that fed). Flies with insufficiently colored abdomen were discarded as poorly fed flies.

### Data analyses

Data analyses were performed using our custom-written R and python programs.

### Statistics

We did not predetermine the sample sizes. The distribution of normality of each group was assessed using Shapiro-Wilk normality test. Parametric tests (unequal variances *t*-test known as Welch’s *t*-test or Dunnett’s test) were used only when distributions were deemed normal; otherwise, non-parametric tests were used (Wilcoxon signed rank test, Wilcoxon rank sum test, or Dunn’s test). All statistical tests were two-sided. We employed Bonferroni correction to correct for two tests, and correction implemented in Dunnett’s or Dunn’s test for more than two tests. p < 0.05 after corrections for multiple comparison was deemed statistically significant. Neither randomization nor blinding was performed for the group allocation during experiments or data analysis. All box plots are generated so that center line indicates median, box limits indicate upper and lower quartiles, and whiskers indicate 1.5x interquartile range. All the statistical results are summarized in Supplementary Data 2.

## DATA AVAILABILITY

Source data for all main text and Supplementary Figures are available via request to the corresponding author.

## CODE AVAILABILITY

Source codes used in the current study are available at https://github.com/mtsuji172/heart_behavior.

## ACKNOWLEDGEMENTS

We thank T. Katsuki (ThorLabs) for technical support in setting up the hardware; members of the Emoto Lab for advice and support. This study was supported by JSPS (KAKENHI 24K18157), JST (ACT-X JPMJAX242D), Yamaguchi Educational and Scholarship Foundation to M.T., AMED-CREST (JP21gm310010), Takeda Science Foundation, Torray Foundation, Naito Foundation, and WPI-IRCN to K.E.

## AUTHOR CONTRIBUTIONS

M.T. designed the experiments. M.T., J. D., Y. U., and A.O. performed the experiments. M.T. analyzed the data. M.T. and K.E. interpreted the results and wrote the paper.

## TABLES

**Supplementary Data 1 List of genotypes and sample sizes used in this study**

**Supplementary Data 2 Statistics of data presented in this study**

## MOVIES

**Supplementary Movie 1** An air puff induces cardiac deceleration and promotes locomotion. A 50 mL/min puff is applied in the middle of the movie, as indicated by the red text that appears on the top left panel of the movie. The bottom left panel shows a behavioral recording, with automated body parts tracking (blue dots and lines) superimposed on the raw data. The top right panel shows the raw heartbeat recording, and the bottom right panel shows the pixel-wise, color-coded optical flow (color-to-direction mapping is shown on the top right of the panel). The playback speed is reduced to 0.25 of the original file.

**Supplementary Movie 2** A train of 10 air puffs induce cardiac deceleration in a Hand>GFP fly. The playback speed is reduced to 0.25 of the original file.

**Supplementary Movie 3** Air puffs activate DA-WED neurons. The left panel shows a schematic of the DA-WED neurons and the actual calcium imaging data (averaged over the total recording), capturing both DA-WED neurons in the right hemisphere. An arrow indicates the neuron shown on the right panel. The right panel shows GCaMP signals at time 0 on the top, and the time-lapse imaging from there during which air puffs were applied.

**Supplementary Movie 4** Optogenetic activation of DA-WED neurons induces cardiac deceleration and promotes locomotion. The playback speed is reduced to 0.25 of the original file.

**Supplementary Movie 5** Optogenetic cardiac deceleration promotes locomotion. The playback speed is reduced to 0.25 of the original file.

