## Supplementary Figures for "Dopaminergic Neurons Linking Threat Processing to Cardiac Modulation and Locomotor Responses": Supplementary Fig. 1.pdf

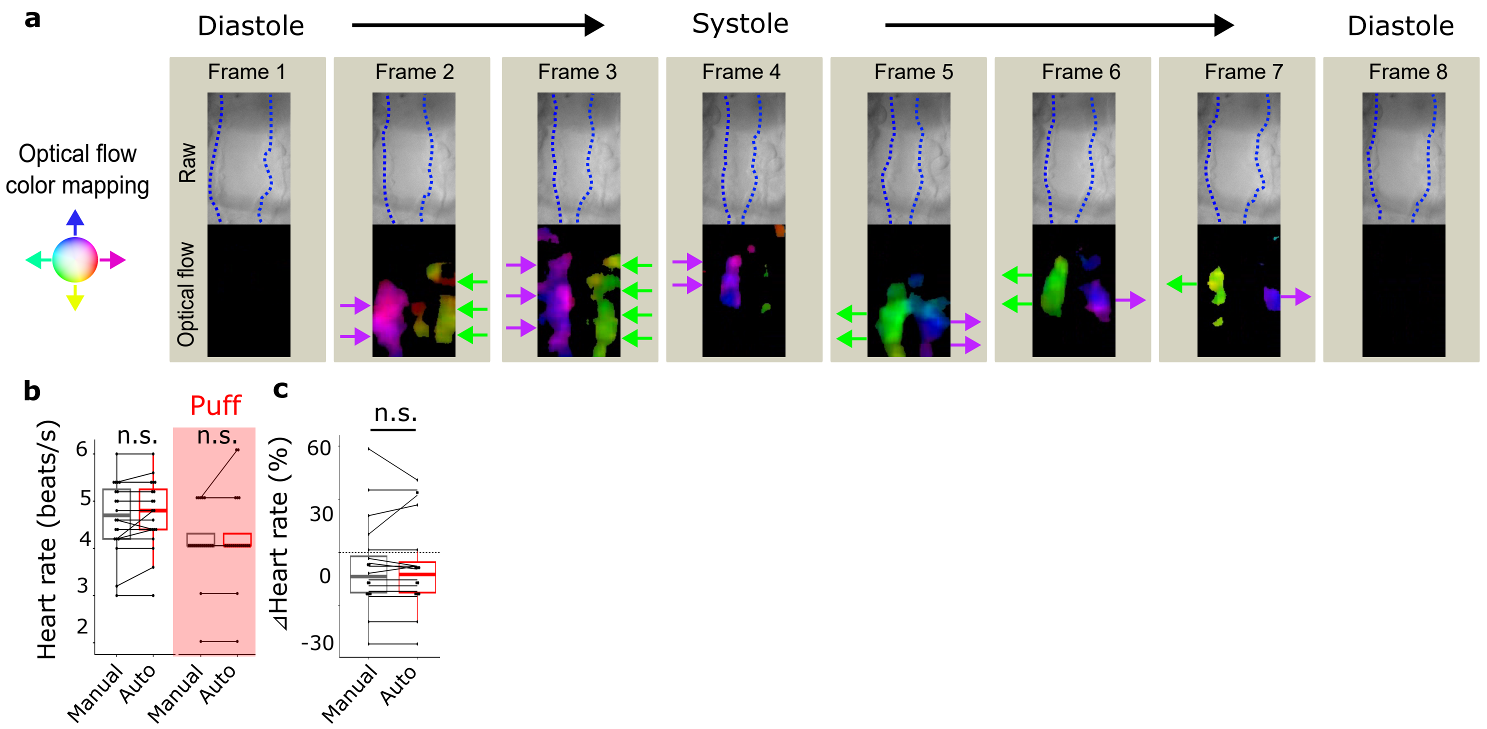


**Supplementary Fig. 1 Heartbeat detection**

**a.** The shown 8 consecutive frames, capture an episode of "diastole -> systole -> diastole" ("Raw", in which blue dotted lines indicate the outline of the conical chamber). Our software estimates the optical flow in these frames ("Optical flow", in which estimated directions are color-coded). Our software reveals a bout of contraction (frames 2 - 4), which is followed by a bout of relaxation (frames 5-8). Arrows indicate the optical flow direction for illustration purposes. **b.** Plot comparing the HR calculated from manual count vs auto-detection, before and during puff application. Each line indicates identical flies. N = 20. n.s.: $p > 0.05$, two-tailed, paired *t*-test followed by Bonferroni correction. **c.** Plot comparing the % change in HR (during vs pre-puff, 10s each) calculated from manual count vs auto-detection. N = 20. n.s.: $p > 0.05$, two-tailed, paired *t*-test.
