## Supplementary Figures for "Dopaminergic Neurons Linking Threat Processing to Cardiac Modulation and Locomotor Responses": Supplementary Fig. 2.pdf

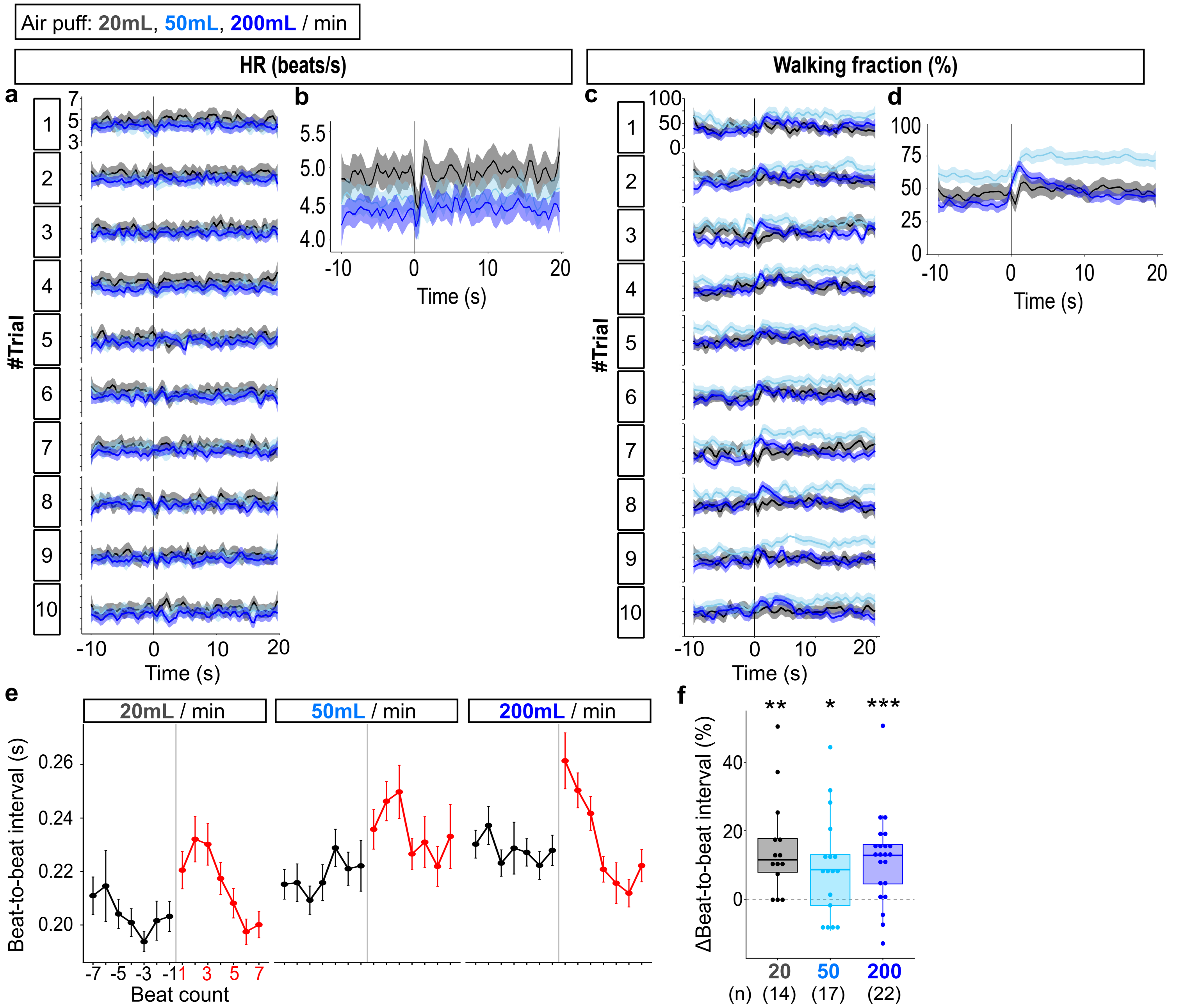


**Supplementary Fig. 2 The effect of air puff on behavior and heartbeat**

**a.** Time course of the HR (beats/s) without averaging over 10 trials. Results of different puff strengths are superimposed (gray: 20mL/min, skyblue: 50mL/min, dark blue: 200mL/min; different fly groups were used for each puff strength). N = 14, 17, 22 for 20mL/min (gray), 50mL/min (sky blue), and 200mL/min (dark blue), respectively. **b.** Time course of the HR averaged over 10 trials for each fly. Results of different puff strengths are superimposed (gray: 20mL/min, skyblue: 50mL/min, dark blue: 200mL/min; different fly groups were used for each puff strength). N = 14, 17, 22 for 20mL/min (gray), 50mL/min (sky blue), and 200mL/min (dark blue), respectively. **c.** Time course of the walking fraction (%) without averaging over 10 trials. Results of different puff strengths are superimposed (gray: 20mL/min, skyblue: 50mL/min, dark blue: 200mL/min; different fly groups were used for each puff strength). N = 14, 17, 22 for 20mL/min (gray), 50mL/min (sky blue), and 200mL/min (dark blue), respectively. **d.** Time course of walking fraction (%) averaged over 10 trials for each fly. Results of different puff strengths are superimposed (gray: 20mL/min, skyblue: 50mL/min, dark blue: 200mL/min; different fly groups were used for each puff strength). N = 14, 17, 22 for 20mL/min (gray), 50mL/min (sky blue), and 200mL/min (dark blue), respectively. **e.** Beat-to-beat interval (in seconds) plotted against the beat count around puff onset (gray vertical line). Dots and bars indicate mean and SEM, respectively. N = 14, 17, 22 for 20, 50, and 200mL/min, respectively. **f.** Change in beat-to-beat interval, calculated for each fly as the % difference between the mean interval during three beats following stimulation onset and the mean interval during three beats preceding stimulation. N = 14, 17, 22 for 20, 50, and 200mL/min, respectively. ***$p < 0.001$, **$p < 0.01$, *$p < 0.05$, Wilcoxon's signed rank test followed by Bonferroni correction.
