## Supplementary Figures for "Dopaminergic Neurons Linking Threat Processing to Cardiac Modulation and Locomotor Responses": Supplementary Fig. 3.pdf

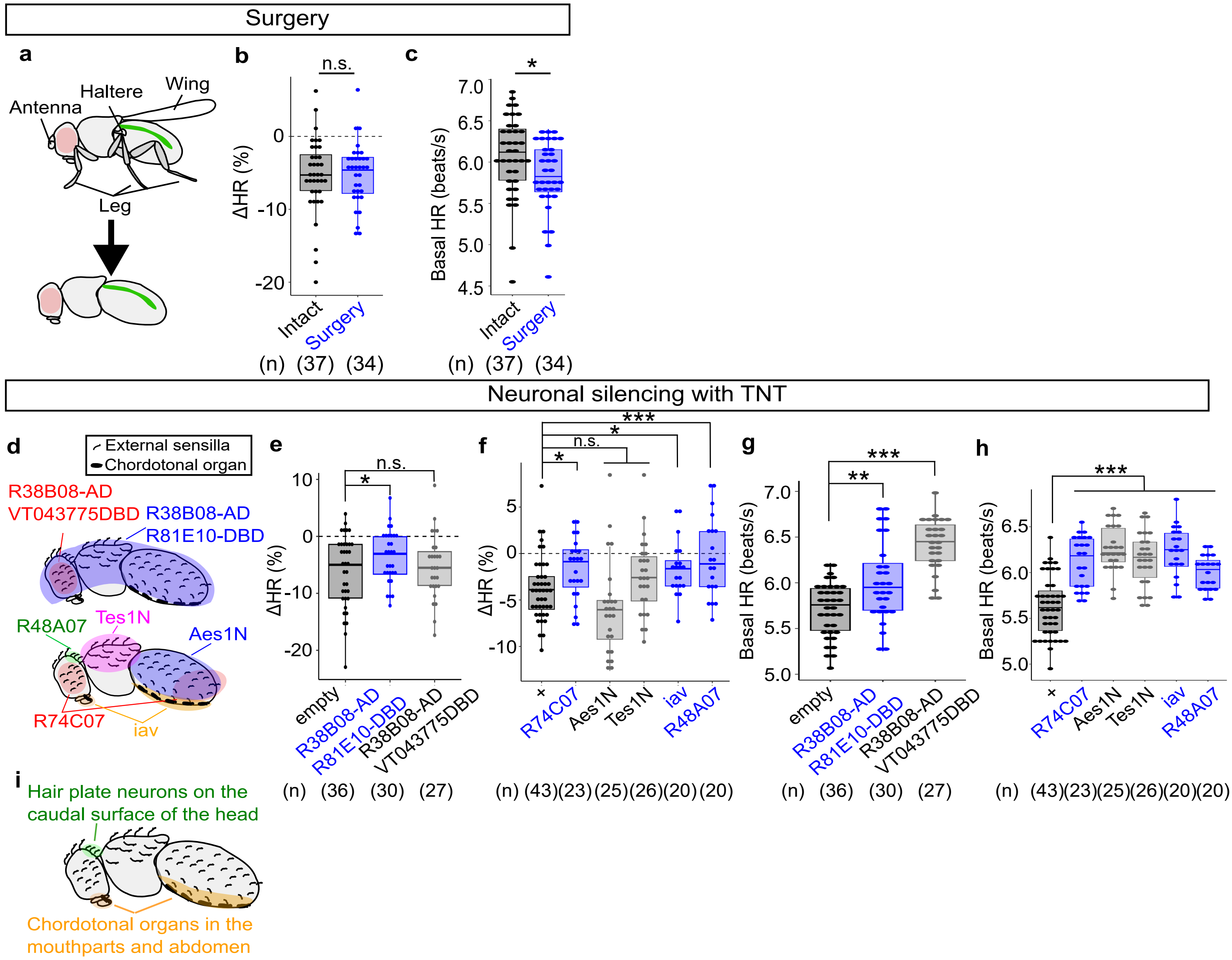


**Supplementary Fig. 3 Somatosensory systems required for puff-induced cardiac deceleration**

**a.** Drosophila possesses multiple mechanosensory organs throughout the body, including legs, haltares, wings, and the body trunk. To first distinguish which of these body parts may be indispensable for puff-induced cardiac deceleration, we surgically removed legs, haltares, wings, and tested puff-induced cardiac deceleration. Green indicates the heart. **b.** Change in HR, calculated for each fly as the % difference between the mean HR during the 10 s stimulation period and the HR HR during the 10 s baseline period preceding stimulation. N = 37, 34, for intact flies, surgery flies, respectively. n.s.: $p > 0.05$, Wilcoxon's rank sum test. **c.** Baseline HR. N = 37, 34, for intact flies, surgery flies, respectively. *$p < 0.05$, two-tailed t-test. **d.** A schematic of the experiment. Different subsets of body hair somatosensors are silenced by expressing TNT using indicated driver lines. (Upper) split GAL4-based experiments. (Lower) GAL-based experiments. **e.** Change in HR, calculated for each fly as the % difference between the mean HR during the 10 s stimulation period and the HR HR during the 10 s baseline period preceding stimulation. N = 36, 30, 27 for "empty", "R38B08-AD R81E10-DBD", "R38B08-AD VT043775-DBD", respectively. *$p < 0.05$, n.s.: $p > 0.05$, Dunnett's test. **f.** Change in HR, calculated for each fly as the % difference between the mean HR during the 10 s stimulation period and the HR HR during the 10 s baseline period preceding stimulation. N = 43, 23, 25, 26, 20, 20 for "+", "R74C07", "Aes1N", "Tes1N", "iav", "R48A07", respectively. ***$p < 0.001$, *$p < 0.05$, n.s.: $p > 0.05$, Dunnett's test. **g.** Baseline HR. N = 36, 30, 27 for "empty", "R38B08-AD R81E10-DBD", "R38B08-AD VT043775-DBD", respectively. ***$p < 0.001$, **$p < 0.01$, Dunnett's test. **h.** Baseline HR. N = 43, 23, 25, 26, 20, 20 for "+", "R74C07", "Aes1N", "Tes1N", "iav", "R48A07", respectively. ***$p < 0.001$, Dunnett's test. **i.** Somatosensory neurons suggested to be partially indispensable for puff-induced cardiac deceleration. When multiple drivers labeled the given neurons, we considered such neurons as indispensable for cardiac deceleration only when those drivers produced consistent results (hence, e.g. external sensilla on the posterior abdomen were excluded based on Aes1N driver's result, despite R74C07 driver's result suggests otherwise; similarly, sensilla on the eye compound labeled by R74C07 driver were excluded by “R38B08-AD VT043775-DBD” split combination’s result.).
