## Supplementary Figures for "Dopaminergic Neurons Linking Threat Processing to Cardiac Modulation and Locomotor Responses": Supplementary Fig. 4.pdf

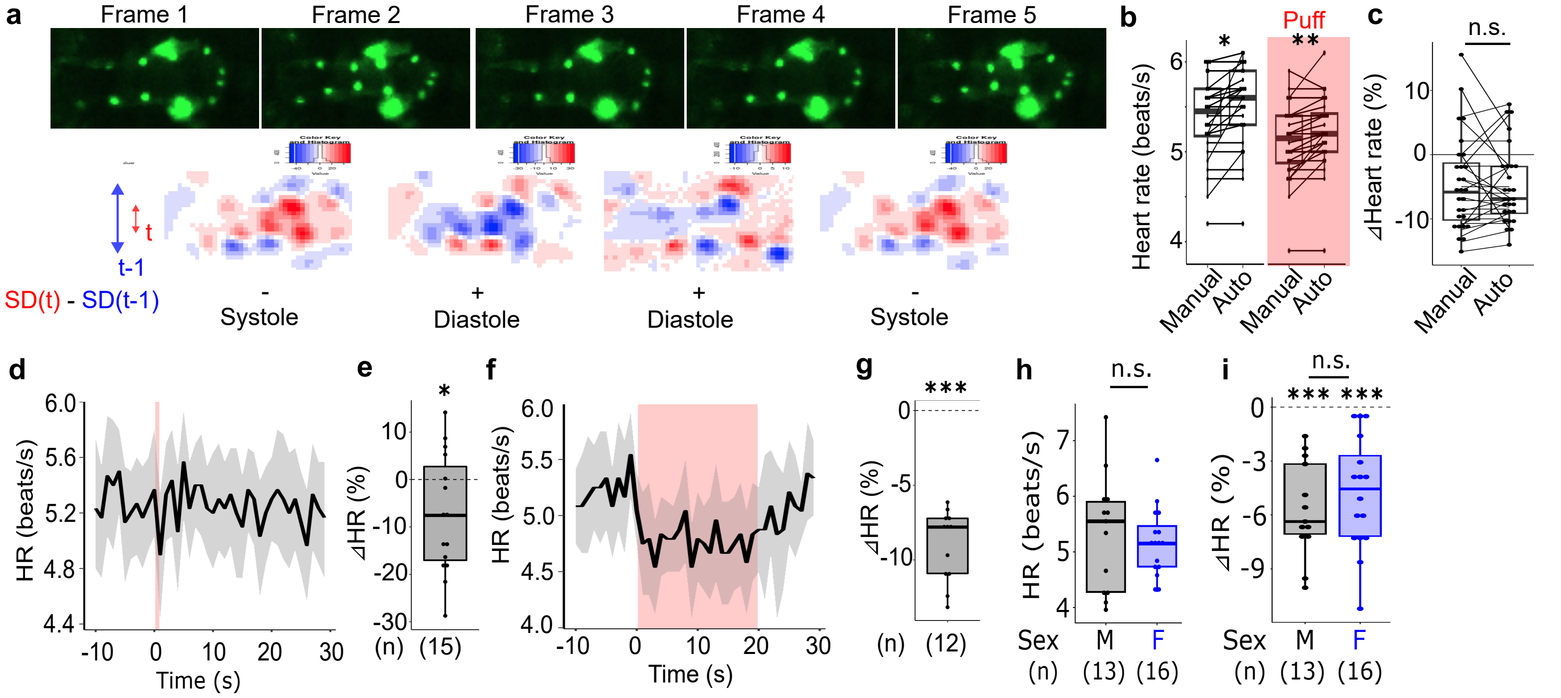


**Supplementary Fig. 4 Heartbeat detection analysis**

**a.** (Upper) example frames capturing the heart tube of a Hand-GFP fly. (Lower) Heat plot showing for each pixel the difference between consecutive frames. Pixels whose value at time t is larger than that at time (t-1) are shown with red; blue for the opposite. Double-headed arrows show the width of the heart for time t and (t-1) for illustrative purposes, and are not meant to be quantitatively precise. "SD(t)" indicates the standard deviation of the y coorinates of red pixels, and "SD(t-1)" of blue pixels. SD(t) is larger when the width of the heart is larger. Thus, "SD(t) - SD(t-1)" indicates systole when < 0, and diastole when > 0. **b.** Plot comparing the HR calculated from manual count vs auto-detection, before and during puff application. Each line indicates identical flies. N = 27. **p < 0.01, *p < 0.05, two-tailed, paired *t*-test followed by Bonferroni correction. **c.** Plot comparing the % change in HR (during vs pre-puff, 10s each) calculated from manual count vs auto-detection. n.s.: p > 0.05, two-tailed *t*-test. **d.** Time course of the HR. A single puff was applied during red-shaded time window. Lines and shaded areas represent means and SEM, respectively. N = 15. **e.** Change in HR, calculated for each fly as the % difference between the mean HR during the 1 s stimulation period and the HR HR during the 10 s baseline period preceding stimulation. N = 15, *p < 0.05, two-tailed *t*-test. **f.** Time course of the HR. A train of 20 puffs were applied during red-shaded time window. Lines and shaded areas represent means and SEM, respectively. N = 12. **g.** Change in HR, calculated for each fly as the % difference between the mean HR during the 20 s stimulation period and the HR HR during the 10 s baseline period preceding stimulation. N = 12, ***p < 0.001, two-tailed *t*-test. **h.** The baseline HR. N = 13, 15 for male, female, respectively. n.s.: p > 0.05, two-tailed *t*-test. **i.** Change in HR, calculated for each fly as the % difference between the mean HR during the 10 s stimulation period and the HR HR during the 10 s baseline period preceding stimulation. N = 13, 15 for male, female, respectively. ***p < 0.001, n.s.: p > 0.05, two-tailed *t*-test. Sex-wise statistics were Bonferroni-corrected.
