## Supplementary Figures for "Dopaminergic Neurons Linking Threat Processing to Cardiac Modulation and Locomotor Responses": Supplementary Fig. 5.pdf

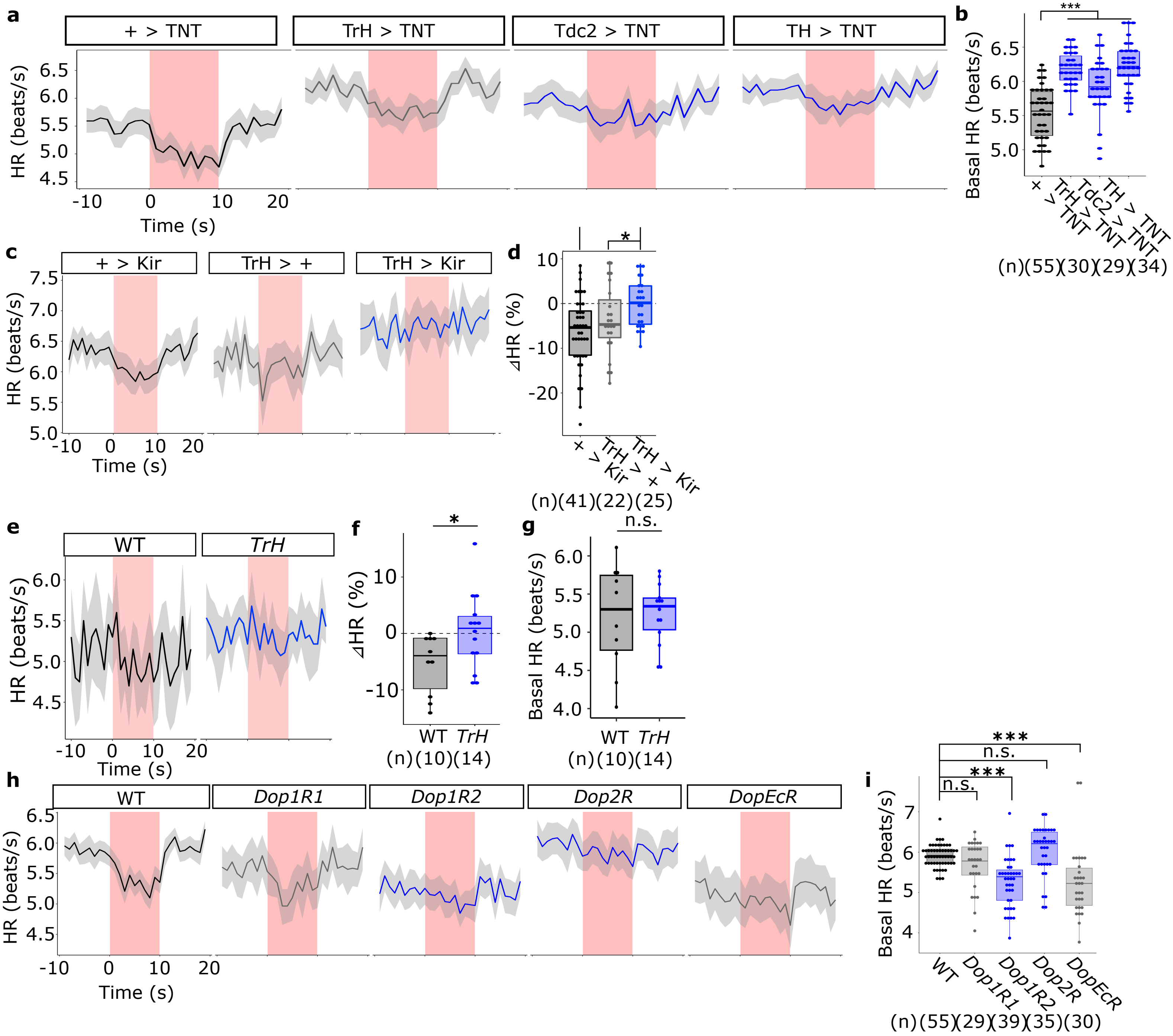


**Supplementary Fig. 5 Roles of monoaminergic systems in puff-induced cardiac deceleration**

**a.** Time courses of the HR in beats per sec. A train of 10 puffs were applied during red-shaded time window. Lines and shaded areas represent means and SEM, respectively. N = 55, 30, 29, 34, for "+", TrH, Tdc2, TH, respectively. **b.** The baseline HR in beats per second. N = 55, 30, 29, 34, for "+", TrH, Tdc2, TH, respectively. ***$p < 0.001$, Dunnett's test. **c.** Time courses of the HR in beats per sec. A train of 10 puffs were applied during red-shaded time window. Lines and shaded areas represent means and SEM, respectively. N = 41, 22, 25, for "+>Kir","TrH>+","TdH>Kir", respectively. **d.** Change in HR, calculated for each fly as the % difference between the mean HR during the 10 s stimulation period and the HR HR during the 10 s baseline period preceding stimulation. N = 41, 22, 25, for "+>Kir","TrH>+","TdH>Kir", respectively. ***$p < 0.001$, *$p < 0.05$, Dunnett's test. **e.** Time courses of the HR in beats per sec. A train of 10 puffs were applied during red-shaded time window. Lines and shaded areas represent means and SEM, respectively. N = 10, 14 for WT, *TrH*, respectively. **f.** Change in HR, calculated for each fly as the % difference between the mean HR during the 10 s stimulation period and the HR during the 10 s baseline period preceding stimulation. N = 10, 14 for WT, *TrH*, respectively. *$p < 0.05$, two-tailed *t*-test. **g.** The baseline HR in beats per second. N = 10, 14 for WT, *TrH*, respectively. n.s.: $p > 0.05$, two-tailed *t*-test. **h.** Time courses of the HR in beats per sec. A train of 10 puffs were applied during red-shaded time window. Lines and shaded areas represent means and SEM, respectively. N = 55, 29, 39, 35, 30, for WT, *Dop1R1*, *Dop1R2*, *Dop2R*, *DopEcR*, respectively. **i.** The baseline HR in beats per second. N = 55, 29, 39, 35, 30, for WT, *Dop1R1*, *Dop1R2*, *Dop2R*, *DopEcR*, respectively. ***$p < 0.001$, n.s.: $p > 0.05$, Dunn's test.
