## Supplementary Figures for "Dopaminergic Neurons Linking Threat Processing to Cardiac Modulation and Locomotor Responses": Supplementary Fig. 6.pdf

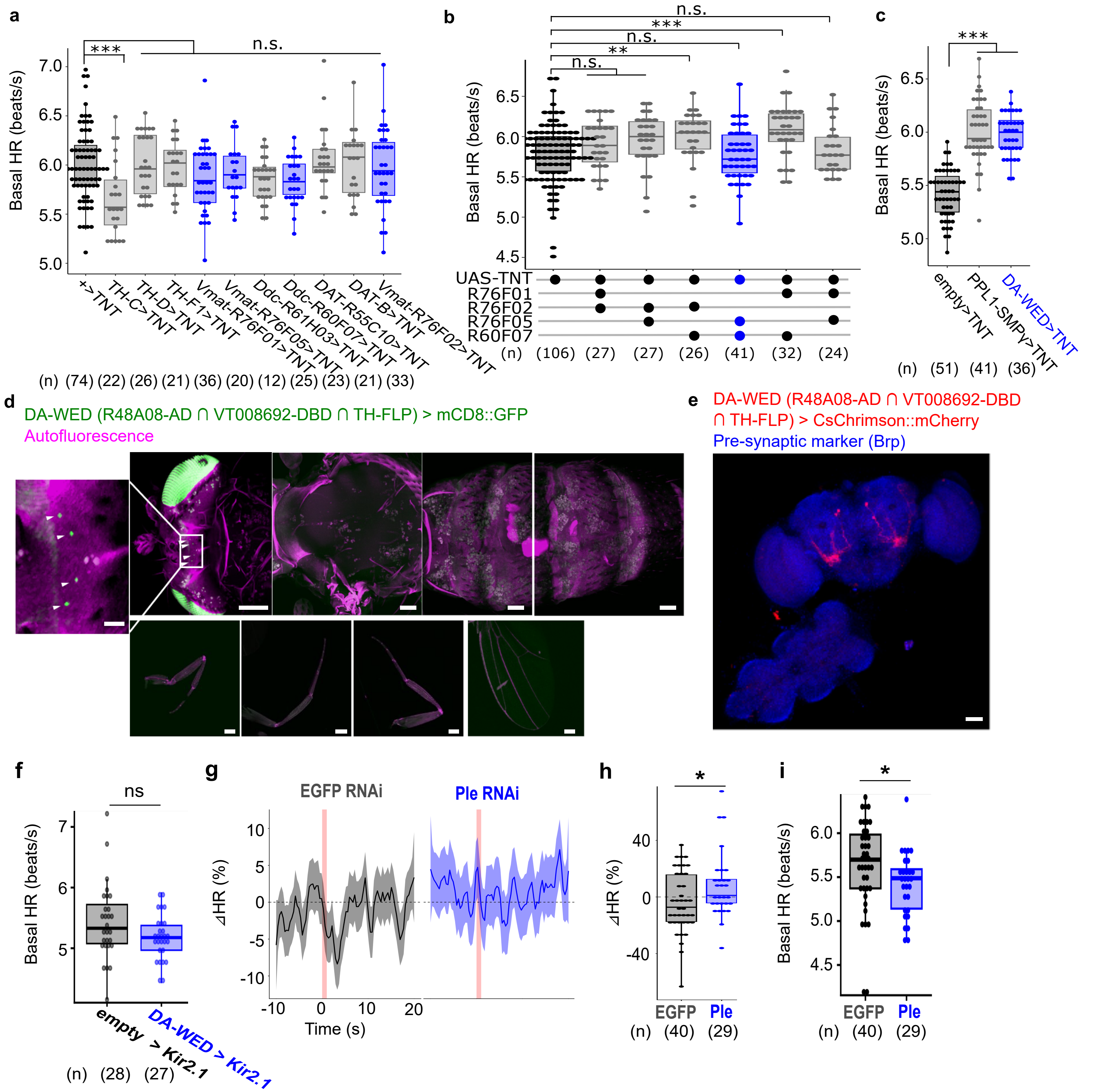


**Supplementary Fig. 6 Silencing different subsets of dopaminergic neurons, DA-WED driver's expression pattern, DA-WED silencing, and ple RNAi**

**a.** The baseline HR in beats per second. N = 74, 22, 26, 21, 36, 20, 12, 25, 23, 21, 33, for "+", TH-C, TH-D, TH-F1, Vmat-R76F01, Vmat-R76F05, Ddc-R61H03, Ddc-R60F07, DAT-R55C10, DAT-B, Vmat-R76F02, respectively. ***$p < 0.05$, n.s.: $p > 0.05$, Dunn's test. **b.** The baseline HR in beats per second. N = 106, 27, 27, 26, 41, 32, 24, for driver-less control, R76F01 ∩ R76F02, R76F02 ∩ R76F05, R76F01 ∩ R76F02, R76F02 ∩ R76F07, R76F05 ∩ R60F07, R76F01 ∩ R60F07, R76F01 ∩ R76F05, respectively.***$p < 0.001$, **$p < 0.01$, n.s.: $p > 0.05$, Dunn's test. **c.** The baseline HR in beats per second. N = 51, 41, 36, for empty, PPL1-SMPγ, DA-WED, respectively. ***$p < 0.001$, Dunn's test. **d.** Cleared tissue of a fly expressing mCD8GFP in DA-WED neurons (R48A08-AD ∩ VT008692-DBD ∩ TH-FLP > mCD8::GFP (green). Autofluorescence of the tissue is visible as magenta. Arrowheads indicate DA-WED neuron's soma. The bottom panels each show, from left to right, a foreleg, midleg, hindleg, and wing. Scale bar: 100 um (inlet: 25 um). **e.** A brain and VNC of R48A08-AD ∩ VT008692-DBD ∩ TH-FLP > CsChrimson::mCherry (red) immunostained with anti-neuropil marker nc82 (blue). Scale bars: 25 um (inlet: 5 um). Similar results were obtained across 7 independent samples. Scale bar: 50 um. **f.** The baseline HR in beats per second. N = 28, 27, for empty>Kir2.1, DA-WED>Kir2.1, respectively. n.s.: $p > 0.05$, two-tailed *t*-test. **g.** Time course of the HR (% change from baseline defined as the average during the 10 s preceding puff onset). An air puff was applied during the red-shaded time window. Lines and shaded areas represent means and SEM, respectively. N = 40, 29, for EGFP-RNAi, ple-RNAi, respectively. **h.** Change in HR, calculated for each fly as the % difference between the mean HR during the 1 s following stimulation onset and the mean HR during the 10 s baseline period preceding stimulation. N = 40, 29, for EGFP-RNAi, ple-RNAi, respectively. *$p < 0.05$, two-tailed *t*-test. **i.** The baseline HR. N = 40, 29, for EGFP-RNAi, Ple-RNAi, respectively. *$p < 0.05$, Wilcoxon' signed rank test.
