## Supplementary Figures for "Dopaminergic Neurons Linking Threat Processing to Cardiac Modulation and Locomotor Responses": Supplementary Fig. 7.pdf

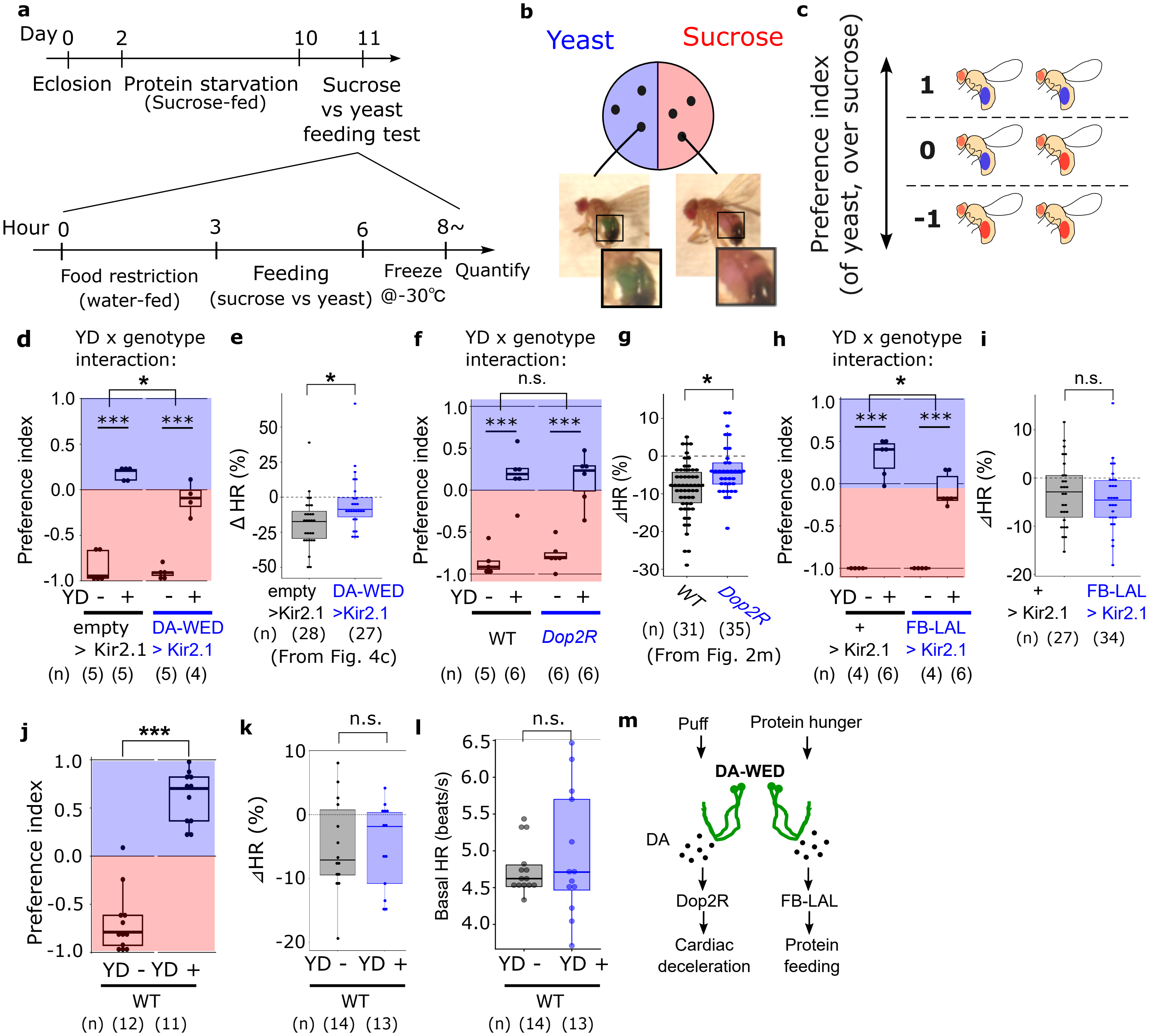


**Supplementary Fig. 7 DA-WED neurons induce cardiac deceleration and protein feeding through distinct mechanisms**
**a.** Time course of the experiment. Flies two days post eclosion were subjected to 8-day protein starvation protocol. Protein preference of the starved flies were then tested the next day. The preference test consisted of 3-hour food restriction, then 3-hour feeding on sucrose (supplemented with red dye) or yeast (supplemented with blue dye). The flies were then frozen and their abdominal color was inspected. **b.** Protein preference test. Flies were transferred into a 35mm petri dish, one half of which contained agarose gel with yeast + blue dye, while the other half containing sucrose + red dye. **c.** Preference index was calculated as (# flies consuming yeast - # flies consuming sucrose) / (total # flies that fed). **d.** Preference index. N = 5, 5, 5, 4 batches (each with ~60 flies) for empty YD-, empty YD+, DA-WED YD-, DA-WED YD+, respectively. ***p < 0.001, *p < 0.05, two-tailed *t*-test. YD- vs YD+ statistics were Bonferroni-corrected. **e.** Change in HR, calculated for each fly as the % difference between the mean HR during the 1 s following stimulation onset and the mean HR during the 10 s baseline period preceding stimulation. N = 28, 27, for empty>Kir2.1, DA-WED>Kir2.1, respectively. *p < 0.05, Two-tailed t-test. Fig.4c is shown here again for comparison with d. **f.** Preference index. N = 5, 6, 6, 6 batches for WT YD-, WT YD+, Dop2R YD-, Dop2R YD+, respectively. *p < 0.05, two-tailed *t*-test. YD- vs YD+ statistics were Bonferroni-corrected. **g.** Change in HR, calculated for each fly as the % difference between the mean HR during the 10 s stimulation period and the HR HR during the 10 s baseline period preceding stimulation. N = 31, 35, for WT, Dop2R, respectively. Fig. 2m is shown here again for comparison with f. *p < 0.05, Dunn's test. **h.** Preference index. N = 4, 6, 4, 6 batches for empty YD-, empty YD+, FB-LAL YD-, FB-LAL YD+, respectively. ***p < 0.001, *p < 0.05, two-tailed *t*-test. YD- vs YD+ statistics were Bonferroni-corrected. **i.** Change in HR, calculated for each fly as the % difference between the mean HR during the 1 s following stimulation onset and the mean HR during the 10 s baseline period preceding stimulation. N = 27, 34, for empty>Kir2.1, FB-LAL>Kir2.1, respectively. n.s.: p > 0.05, two-tailed *t*-test. **j.** Preference index. N = 12, 11 batches for YD-, YD+, respectively. ***p < 0.001, two-tailed *t*-test. **k.** Change in HR, calculated for each fly as the % difference between the mean HR during the 1 s following stimulation onset and the mean HR during the 10 s baseline period preceding stimulation. N = 14, 13, for YD-, YD+, respectively. n.s.: p > 0.05, two-tailed *t*-test. **l.** Baseline HR in beats per sec. N = 14, 13, for YD-, YD+, respectively. n.s.: p > 0.05, Wilcoxon's rank sum test. **m.** model.
