## Supplementary Figures for "Dopaminergic Neurons Linking Threat Processing to Cardiac Modulation and Locomotor Responses": Supplementary Fig. 8.pdf

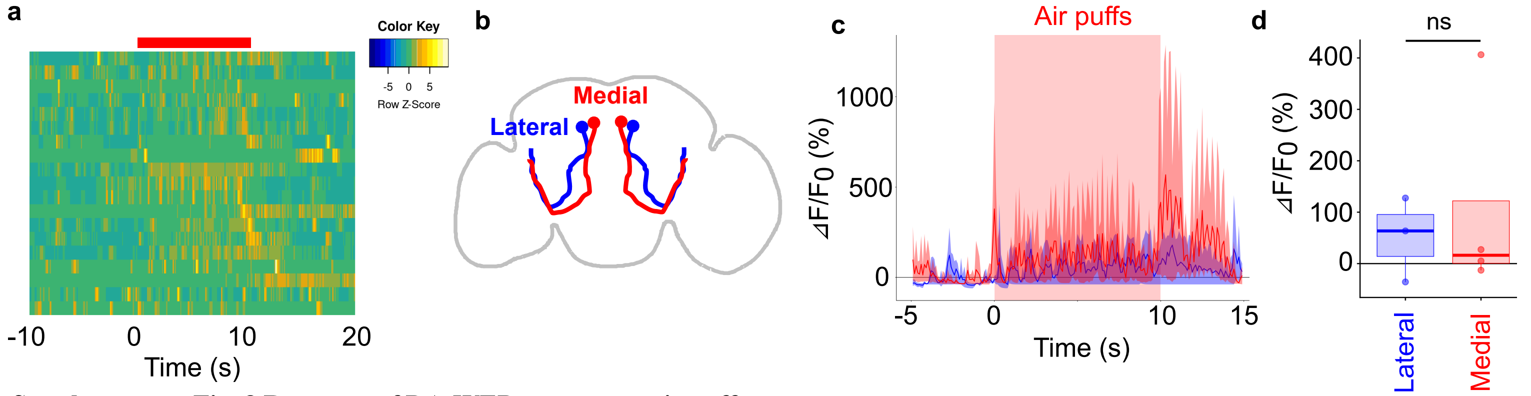


**Supplementary Fig. 8 Response of DA-WED neurons to air puffs**

**a.** Rows depict the color-coded activity of a cell (z score). Red bar indicates the time window during which air puffs were applied. N = 19 cells from 6 flies. **b.** A cartoon showing medial and lateral DA-WED neurons. **c.** Time course of $\Delta F/F_0$. Puffs were applied at 1Hz for 10 times (over 10 s). **d.** $\Delta F/F_0$ averaged over 10s of puff application. When multiple neurons were recorded from the same fly (max 2 neurons per fly), results were averaged so that each dot represents one fly. Neurons for which medial/lateral location could not be distinguished are excluded from this analysis. N = 3, 4 neurons for lateral and medial, respectively. Two-tailed *t*-test. ns: $p > 0.05$.
