## Supplementary Figures for "Dopaminergic Neurons Linking Threat Processing to Cardiac Modulation and Locomotor Responses": Supplementary Fig. 9.pdf

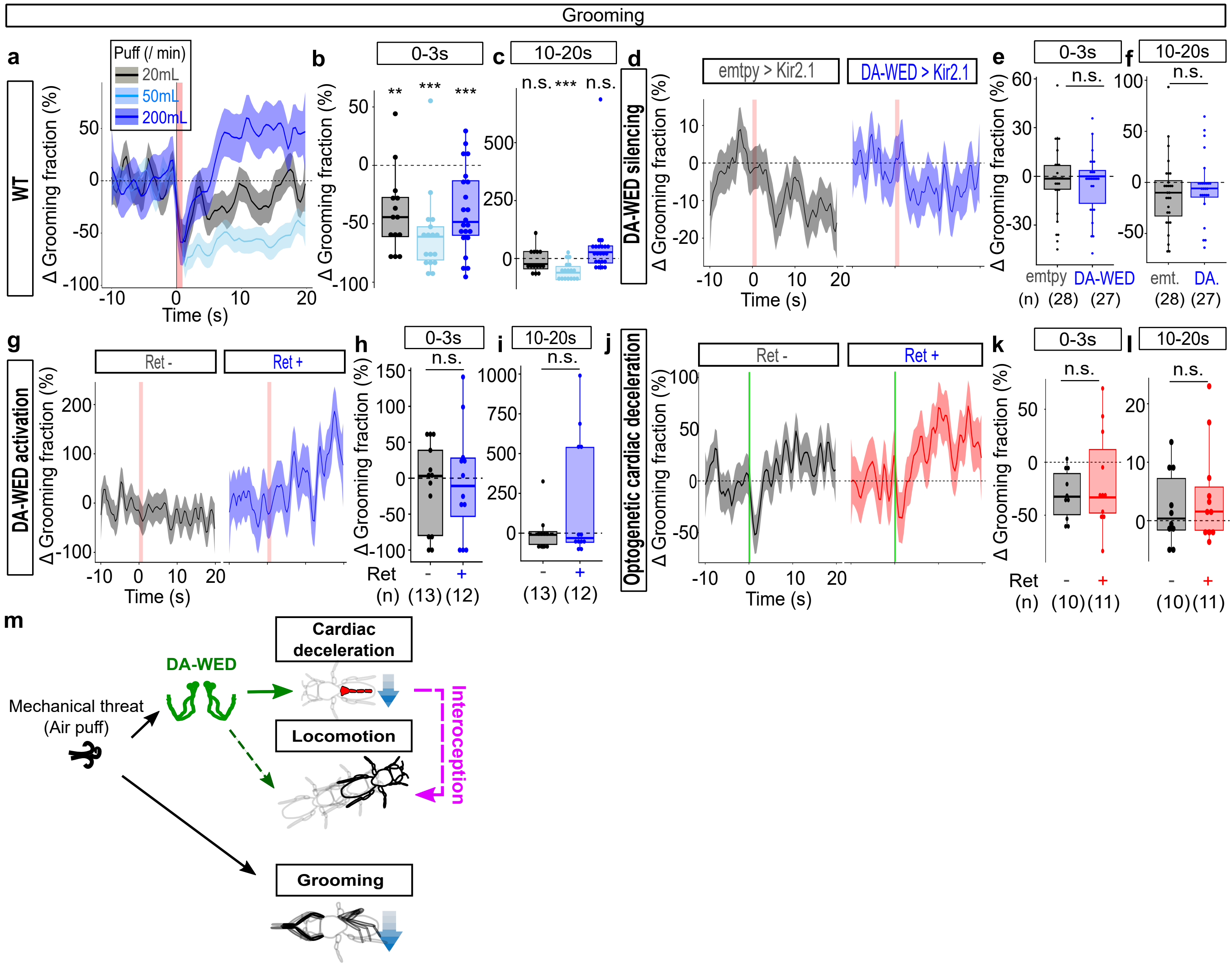


**Supplementary Fig. 9 The effects of air puff, DA-WED manipulation, and optogenetic cardiac deceleration on grooming behavior.**

**a.** Time course of the grooming fraction (% change from baseline defined as the mean fraction during the 10 s preceding puff onset). An air puff was applied during the red-shaded time window. Lines and shaded areas represent means and SEM, respectively. N = 14, 17, 22 for 20 mL/min (gray), 50 mL/min (sky blue), and 200 mL/min (dark blue), respectively. **b, c.** Change in grooming fraction, calculated for each fly as the % difference between the mean fraction during the 3 s (b) or 10 - 20s (c) following stimulation onset and the mean fraction during the 10 s baseline period preceding stimulation. N = 14, 17, 22 for 20 mL/min (gray), 50 mL/min (sky blue), and 200 mL/min (dark blue), respectively. ***$p < 0.001$, **$p < 0.01$, Wilcoxon's signed rank test followed by Bonfferoni correction. **d.** Time course of the grooming fraction (% change from baseline defined as the mean fraction during the 10 s preceding puff onset). An air puff was applied during the red-shaded time window. Lines and shaded areas represent means and SEM, respectively. N = 28, 27, for empty>Kir2.1, DA-WED>Kir2.1, respectively. **e, f.** Change in grooming fraction, calculated for each fly as the % difference between the mean fraction during the 3 s (e) or 10 - 20s (f) following stimulation onset and the mean fraction during the 10 s baseline period preceding stimulation. N = 28, 27, for empty>Kir2.1, DA-WED>Kir2.1, respectively. n.s.: $p > 0.05$, Two-tailed $t$-test. **g.** Time course of the grooming fraction (% change from baseline defined as the mean fraction during the 10 s preceding puff onset). Light was shined during the red-shaded time window. Lines and shaded areas represent means and SEM, respectively. N = 13, 12, for ret -, ret +, respectively. **h, i.** Change in grooming fraction, calculated for each fly as the % difference between the mean fraction during the 3 s (h) or 10 - 20s (i) following stimulation onset and the mean fraction during the 10 s baseline period preceding stimulation. N = 13, 12, for ret -, ret +, respectively. n.s.: $p > 0.05$, Two-tailed $t$-test. **j.** Time course of the grooming fraction (% change from baseline defined as the mean fraction during the 10 s preceding puff onset). Light was shined during the green-shaded time window. Lines and shaded areas represent means and SEM, respectively. N = 10, 11, for ret -, ret +, respectively. **k, l.** Change in grooming fraction, calculated for each fly as the % difference between the mean fraction during the 3 s (k) or 10 - 20s (l) following stimulation onset and the mean fraction during the 10 s baseline period preceding stimulation. N = 10, 11, for ret -, ret +, respectively. n.s.: $p > 0.05$, two-tailed $t$-test. **m.** Model. Air puff promotes locomotion while suppressing grooming. Mechanistically, air puff activates DA-WED neurons. DA-WED neurons drive not only cardiac deceleration but locomotion, potentially through a feedback from cardiac deceleration (dashed arrows). Such a DA-WED-based mechanism fails to influence grooming.
