## Supplementary Figures for "Dopaminergic Neurons Linking Threat Processing to Cardiac Modulation and Locomotor Responses": Supplementary Fig. 10.pdf

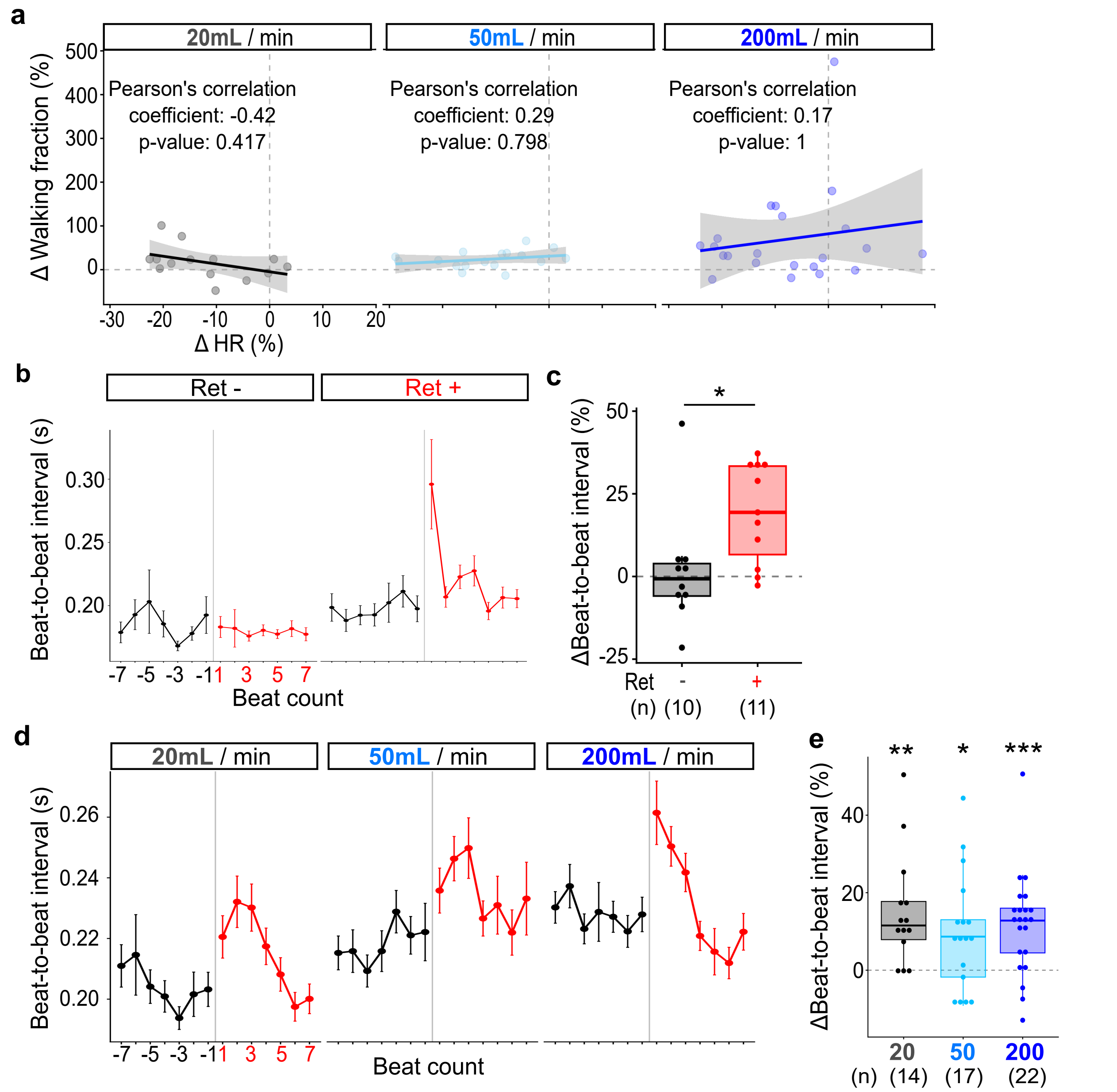


(From Supplementary Fig. 2e)

(From Supplementary Fig. 2f)

**Supplementary Fig. 10 The effects of optogenetic cardiac deceleration on beat-to-beat interval**

**a.** Correlations between changes in walking fraction and HR. N = 14, 17, 22 for 20, 50, and 200mL/min, respectively. Pearson's correlation coefficient, followed by Bonferroni correction. **b.** Beat-to-beat interval (in seconds) plotted against the beat count around puff onset (gray vertical line). Dots and bars indicate mean and SEM, respectively. N = 11, 10 for ret -, +, respectively. **c.** Change in beat-to-beat interval, calculated for each fly as the % difference between the mean interval during three beats following stimulation onset and the mean interval during three beats preceding stimulation. N = 11, 10 for ret -, +, respectively. *$p < 0.05$, Wilcoxon's signed rank test. **d.** Beat-to-beat interval (in seconds) plotted against the beat count around puff onset (gray vertical line). Dots and bars indicate mean and SEM, respectively. N = 14, 17, 22 for 20, 50, and 200mL/min, respectively. Supplementary Fig. 2e is shown here again for comparison. **e.** Change in beat-to-beat interval, calculated for each fly as the % difference between the mean interval during three beats following stimulation onset and the mean interval during three beats preceding stimulation. N = 14, 17, 22 for 20, 50, and 200 mL/min, respectively. ***$p < 0.001$, **$p < 0.01$, *$p < 0.05$, Wilcoxon's signed rank test followed by Bonfferoni correction. Supplementary Fig. 2f is shown here again for comparison.
